# Developmentally inspired bioprinting of nascent multicellular human heart tissue through *in situ* differentiation and morphogenesis of iPSCs

**DOI:** 10.64898/2026.01.13.699298

**Authors:** Ankita Pramanick, Daniel Kelly, Abhay Pandit, Andrew C. Daly

## Abstract

Traditional heart tissue bioprinting typically relies on using human induced pluripotent stem cell (iPSC)-derived cardiomyocytes that are pre-differentiated in 2D culture. This approach differs fundamentally from embryonic heart development, where mesodermal progenitors differentiate into cardiomyocytes within 3D, matrix-rich, and shape-morphing microenvironments. Here, we introduce a novel developmentally inspired approach that enables *in situ* mesodermal and cardiac differentiation of iPSCs within bioprinted, shape-morphing pluripotent tissues. Using embedded bioprinting, Matrigel bioinks with high-density iPSC suspensions were deposited into granular support hydrogels to generate pluripotent tissue constructs with defined architectures. These constructs exhibited shape-morphing behaviour, tunable by modulating the support bath viscoelasticity. Support bath mechanics also regulated iPSC fate, with softer formulations reducing spontaneous differentiation. Building on this, mesodermal induction and cardiogenesis were directly driven within the morphing constructs via temporal WNT pathway modulation, resulting in multicellular cardiac tissues in which cardiomyocytes, fibroblasts, and endothelial cells co-emerge from a common progenitor pool. Importantly, these nascent tissues underwent structural maturation, with immunofluorescence and gene expression profiling revealing cardiac progenitors alongside maturing cardiomyocytes. Together, these findings highlight the potential for a new paradigm in biofabrication focused on printing pluripotent organ rudiments that recapitulate key aspects of embryonic development and support progressive tissue maturation.

## 1. Introduction

Bioprinting has emerged as a powerful approach for fabricating human cardiac tissues by combining induced pluripotent stem cells (iPSCs), bioinks, and additive manufacturing ^1–4^. Most cardiac bioprinting strategies load bioinks with iPSC-derived cardiomyocytes (iPSC-CMs) that have been pre-differentiated in two-dimensional (2D) culture and then printed into three-dimensional (3D) constructs with defined geometries. This paradigm has yielded promising results, producing contractile cardiac tissue constructs that exhibit spontaneous and synchronous contractions for several months ^2^. Additionally, co-culturing iPSC-CMs with supporting fibroblasts and endothelial cells can enhance tissue organisation and functional maturation ^3,5^. Despite these advantages, pre-differentiation strategies differ significantly from the development of the embryonic heart. For example, iPSC-CMs are generated in 2D microenvironments, then dissociated and re-embedded into bioinks, steps that disrupt their native cell-matrix and cell-cell interactions. This approach bypasses the natural developmental trajectory from pluripotency to mesodermal progenitors to cardiomyocytes, which occurs within a 3D, matrix-rich, and mechanically evolving microenvironment. Furthermore, supporting cells, such as fibroblasts and endothelial cells, are typically generated using separate differentiation protocols, limiting the intercellular signalling that accompanies developmental co-emergence.

Parallel progress in cardiac organoid engineering has shown that ‘*in situ*’ differentiation of iPSCs can generate multicellular tissues that recapitulate key features of early heart development ^6,7^. In these systems, iPSCs differentiate into mesoderm progenitors, followed by cardiac lineage specification, leading to the co-emergence of multiple cardiac cell types, including cardiomyocytes, endothelial cells, and fibroblasts ^6–8^. Importantly, these self-organising organoids or cardioids also exhibit spatial patterning of myocardial, endocardial, and epicardial cells and can form cavity-containing structures reminiscent of nascent heart chambers ^6^. Employing such biomimetic ‘*in situ*’ differentiation approaches for cardiac bioprinting holds great promise, as printing affords greater geometrical control and scalability compared to organoid systems. One study has demonstrated the successful *in situ* differentiation of bioprinted iPSCs into iPSC-CMs within GelMA/collagen-based bioinks to produce contractile heart tissues ^9^. However, iPSC-CM differentiation was localised to the peripheral regions of the constructs, with the central areas becoming acellular by the end of the culture period. This may have been due to the use of a covalently crosslinked bioink that affected iPSC cellular interactions, viability, and self-organisation.

Developing bioinks that support iPSC viability and self-organisation during the early stages of culture is a critical consideration for *in situ* differentiation strategies. For example, it is well established that iPSCs and embryonic stem cells require cell-cell contact and aggregation for survival ^10–13^. The bioink must be capable of promoting intercellular interactions following encapsulation, and viscoelastic matrices have been shown to enhance iPSC viability, proliferation, and morphogenesis ^14,15^. Beyond cell-scale behaviours, tissue-scale morphogenesis is also a critical consideration. During embryogenesis, cardiac differentiation coincides with morphogenetic processes, such as heart tube formation and looping ^16–18^. Inspired by this, our prior work has shown that cell-mediated shape-morphing can enhance the structural and functional properties of bioprinted heart tissues ^5^. In these studies, the contractile forces generated by fibroblasts induced programmable shape-morphing in collagen-based bioinks, with the resulting internal stresses promoting cardiomyocyte alignment and sarcomere maturation. However, because these constructs were bioprinted using pre-differentiated iPSC-CMs and adult cardiac fibroblasts, morphogenetic shape changes were decoupled from the early differentiation transitions.

Here, we present a developmentally inspired bioprinting strategy that combines shape-morphing with *in situ* cardiac lineage specification. Using embedded bioprinting, we deposited Matrigel bioinks containing high-density suspensions of iPSCs (150 million cells mL^−1^) into soft granular support hydrogels that preserved the print geometry while permitting cell-driven morphogenesis. We first sought to identify support bath formulations that maintain iPSC pluripotency while minimising spontaneous differentiation. We aimed to direct mesodermal differentiation and cardiogenesis within these shape-morphing constructs by temporally modulating the WNT pathway. We hypothesised that synchronising differentiation with morphogenesis would produce nascent cardiac tissue capable of endogenous structural and functional maturation.

## 2. Main

### 2.1 Bioprinting iPSCs in granular support hydrogels

First, we established a method for bioprinting 3D constructs composed of iPSCs. To achieve this, iPSCs were suspended at a high-density in a Matrigel-based bioink and extruded into agarose granular support baths using an embedded bioprinting technique (Figure 1a, i and ii). The Matrigel bioink was maintained at 6-8°C in a non-gelled state to enable extrusion (Supplementary Figure 1). The shear-thinning and self-healing properties of the granular support hydrogel facilitated precise deposition of the low-viscosity bioink, allowing for the fabrication of complex 3D architectures (Figure 1a i, ii). Additionally, the granular support provided mechanical stability to preserve construct geometry during the early stages of culture. Confocal imaging 24 h post-printing confirmed that iPSCs within the printed constructs expressed the pluripotency markers OCT4 and SOX2, as well as the proliferation marker Ki67, indicating the maintenance of stemness and active cell division (Figure 1a ii). After demonstrating that undifferentiated iPSCs could be bioprinted into 3D constructs within granular support hydrogels while retaining pluripotency, the next objective was to investigate whether these cells could undergo *in situ* mesodermal induction and differentiate into cardiomyocytes directly within the bioink (Figure 1b). To optimise the efficiency of this *in situ* differentiation process, we next examined how the composition of both the bioink and the granular support hydrogel could impact iPSC viability and differentiation post-printing.

**Figure 1.**
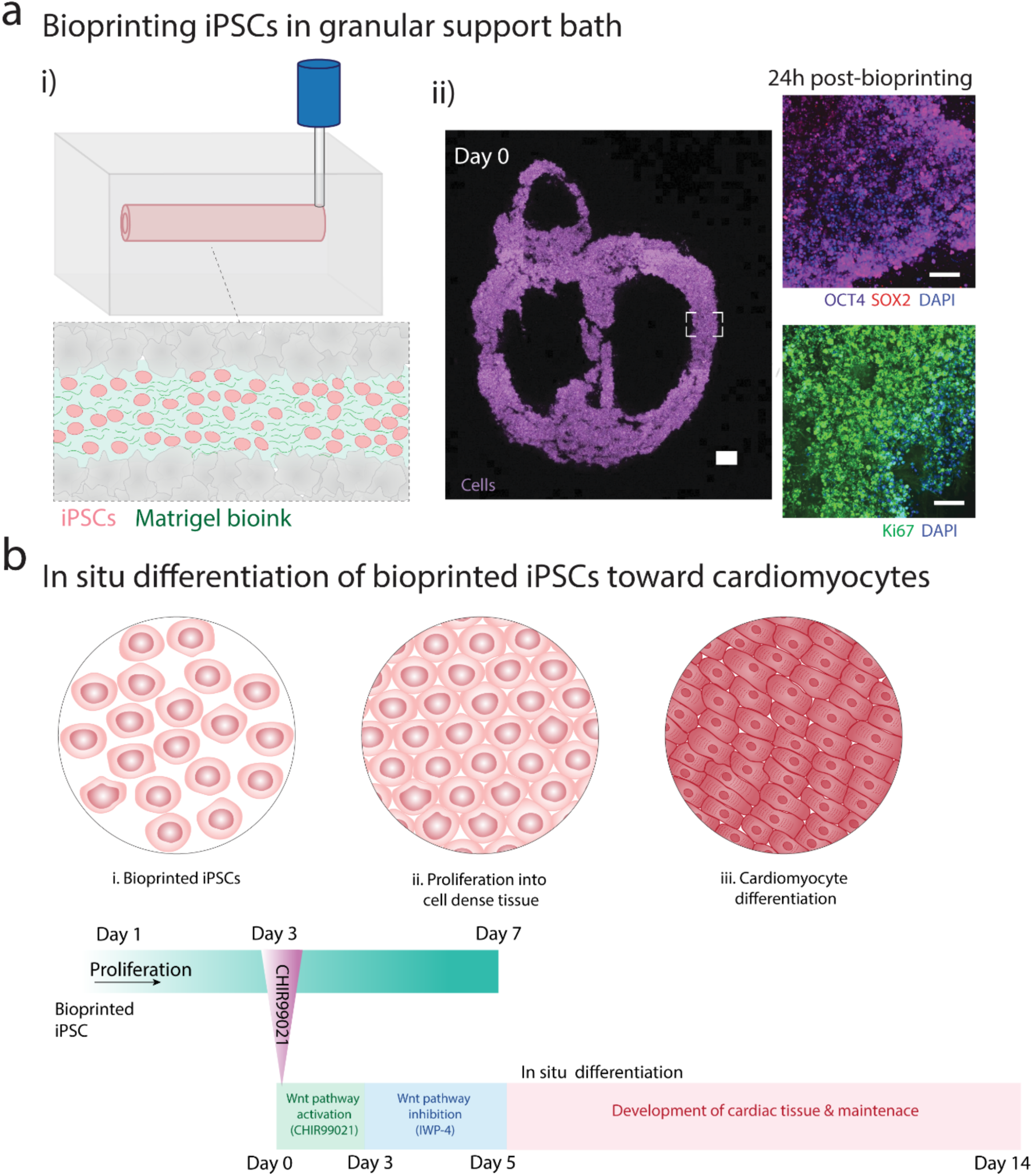
Embedded bioprinting of iPSCs in a granular bath supporting proliferation and *in situ* differentiation: **a)** i) Schematic representation of iPSC bioprinting in a granular support bath. ii) Confocal image of the chambered heart cross-section bioprinted in an agarose support bath (left, scale 1000 µm) and immunofluorescence staining of pluripotency (OCT4, SOX2) and proliferation (Ki67) markers after bioprinting (scale 100 µm). **b)** Schematic illustration of bioprinted iPSCs undergoing proliferation within the bioink prior to differentiation into cardiomyocytes via WNT pathway modulation using the modified GiWi protocol ^19^.

Bioprinting iPSCs as single-cell suspensions presents challenges due to their sensitivity and dependence on cell-cell interactions for survival ^10^. These interactions are essential for maintaining pluripotency and preventing spontaneous differentiation. In addition, external stresses, such as shear forces during extrusion or unfavourable bioink properties, can compromise viability and pluripotency. To mitigate these effects, high-density iPSC suspensions (15-150 million cells mL^−1^) were used for bioprinting to enhance cell-cell interactions. Constructs bioprinted with the highest cell density (150 million cells mL^−1^) maintained approximately 90% viability for up to 7 days of culture (Figure 2a, i and ii, Supplementary Figure 2a). Although lower cell densities (15 and 50 million cells mL^−1^) exhibited slightly higher viability (>95%), these constructs were mechanically weaker and prone to structural failure during culture (Supplementary Figure 2b i, ii, iii). In contrast, constructs bioprinted at 150 million cells mL^−1^ displayed consistent structural integrity, likely due to enhanced cell-cell adhesion. Confocal imaging confirmed the formation of densely packed cells at higher printing densities (Figure 1a, ii). Furthermore, Ki67 staining on day 1 indicated active proliferation of the printed iPSCs (Figure 1a, ii), which likely further enhanced cell-cell adhesion and construct stability.

**Figure 2.**
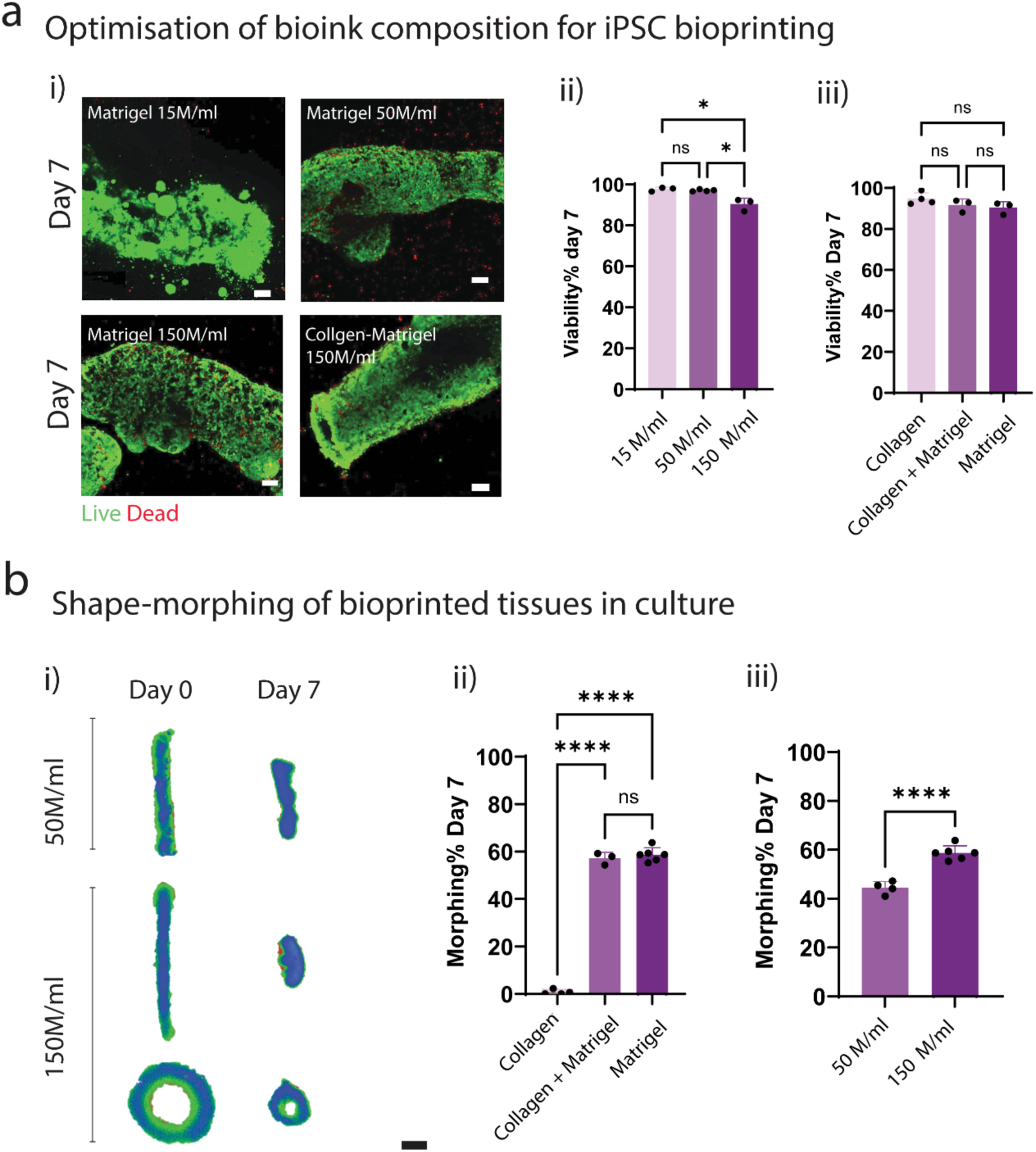
Evaluation of cell viability and tissue shape-morphing of bioprinted constructs: **a)** i) Confocal images indicating the viability within the bioprinted iPSC tissues (scale 100 µm) and ii), iii) quantitative analysis demonstrating the percentage of live cell area as a function of cell density (15, 50, 150 million cells/ml) and ink composition (collagen, collagen + Matrigel, Matrigel), respectively, on day 7. **b)** Bioprinted constructs exhibited tissue shape-morphing during the culture period, as evidenced by i) brightfield images showing the change in tissue dimension over 7 days (scale 1 mm). Quantitative analysis of shape-morphing as a function of ii) bioink composition and iii) cell density. (n = 3 to 6 biological replicates, unpaired t-test for two groups and one-way ANOVA with Tukey’s multiple comparison test for more than two groups, where ns denotes not significant, * denotes p < 0.05, and **** denotes p < 0.0001).

The impact of bioink composition on iPSC behaviour was assessed, focusing on Matrigel and collagen because of their known capacity to support cell-extracellular matrix (ECM) interactions. Matrigel is a well-established matrix for iPSC culture ^20,21^, and collagen hydrogels have also been used for 3D culture of iPSCs ^22,23^. Three formulations were examined: collagen type I alone (0.96 mg mL^−1^), Matrigel alone (3.2-4.4 mg mL^−1^), and a mixed bioink containing both collagen (0.6 mg mL^−1^) and Matrigel (2-2.75 mg mL^−1^). The printed constructs gradually stabilised or stiffened during culture within the support hydrogel, likely due to the formation of cell-cell adhesions and/or nascent ECM production. The printed constructs were cultured within the support hydrogels for 3 days until they established sufficient structural integrity to be removed via the gradual dilution of the support bath with culture media. All bioink formulations supported high cell viability (≥ 90%), with no significant differences observed between them (Figure 2a, iii). However, live/dead staining revealed that the iPSC morphology varied depending on the bioink composition. For example, iPSCs encapsulated in collagen-only bioinks remained largely dispersed as single cells (Supplementary Figure 2a, i), whereas both Matrigel-only and collagen-Matrigel bioinks promoted cell aggregation and intercellular connectivity (Supplementary Figure 2a, i). This enhanced cell aggregation is likely because Matrigel contains proteins such as laminin, collagen IV, and entactin, which support cell adhesion and survival through integrin signalling pathways ^20,21^.

Cell viability was further compared between iPSCs bioprinted as single-cell suspensions and those printed as pre-formed embryoid bodies (EBs) within the Matrigel bioink. The EB culture format is commonly used for iPSC culture as it promotes cell-cell contact and, in the context of extrusion bioprinting, may protect iPSCs from shear stress during extrusion. Despite these potential advantages, both bioprinting approaches using single-cell suspensions or EBs achieved similar high cell viabilities of approximately 90% (Supplementary Figure 2a, i, ii). By day 7, constructs printed from single-cell suspensions exhibited robust cell-cell connectivity (Figure 2a, i), indicating that the iPSCs had self-assembled into geometrically defined 3D structures with dense cellular packing comparable to that of EBs. These results demonstrate that high-density iPSC suspensions in Matrigel bioinks can rapidly establish intercellular connectivity following printing, generating cell-dense, structurally stable, pluripotent tissue constructs. Based on these results, the Matrigel bioink with a cell density of 150 million mL^−1^ was selected for further studies, unless otherwise stated.

### 2.2 Shape-morphing of bioprinted iPSC constructs in granular support hydrogels

Granular support hydrogels provide a soft viscoelastic environment that can accommodate the shape-morphing of bioprinted constructs. For example, our previous work demonstrated that fibroblast-generated contraction forces can drive the shape-morphing of collagen hydrogels in granular supports ^5^. Similarly, in the present study, bioprinted iPSC-containing constructs underwent significant shrinkage and shape-morphing during culture (Figure 2b). These constructs were printed in a soft agarose granular support hydrogel (storage modulus ≈ 105 Pa), which provided structural support during the early stages of culture while allowing morphogenetic deformations. The support bath was gradually diluted and removed by day 3, once the constructs had developed sufficient structural integrity. The magnitude of shape-morphing was influenced by both the bioink composition and cell density (Figure 2b). For example, constructs bioprinted with Matrigel or collagen-Matrigel bioinks underwent greater shape-morphing than those printed with collagen alone, which remained largely static (Figure 2b, ii). Additionally, increasing the iPSC density from 50 to 150 million cells mL^−1^ significantly enhanced shape-morphing in Matrigel-based constructs (Figure 2b, iii). In 2D culture, iPSC colonies have been shown to maintain their morphology and compactness through contractile actin fences and focal adhesions ^24^. These contractile actin fences exert Rho-ROCK-myosin mechanical stresses to maintain colony morphology and compaction. The shape-morphing observed in our bioprinted constructs may be driven by similar mechanisms, whereby iPSCs contract the Matrigel matrix to form intercellular adhesions and densely packed microarchitectures. The constructs bioprinted with both beam– and ring-shaped geometries displayed comparable degrees of shape-morphogenesis (Supplementary Figure 2c, i, ii). In both ring-shaped and beam constructs, morphing increased significantly over time from ∼30-38% on day 3 to ∼ 55-57% by day 7 (Supplementary Figure 2c, iii). It should be noted that some loss of the outer cell and bioink layers occurred within the initial 72 h, but overall, the constructs maintained the desired bioprinted structure (e.g. beam or ring).

To assess how the surrounding mechanical environment affected shape-morphing, constructs were bioprinted and cultured in granular support baths with varying microgel packing densities: high (100%), medium (80%), and low (60%). All formulations exhibited yield stress behaviour, transitioning from elastic to fluid-like states at shear strains of approximately 0.1% (Figure 3a, i, ii). As the packing density decreased from 100% to 60%, the storage modulus decreased from approximately 501 to 105 Pa (Figure 3a, i and ii). The constructs were cultured in the support bath for 7 days (without dilution or removal of the support). As anticipated, the extent of shape-morphing was influenced by the relative elasticity of the support bath. Beam constructs cultured in the stiffest bath (100% packing) underwent the least shape-morphing (13%), whereas constructs cultured in the softest bath (60% packing) underwent significantly greater shape-morphing (65%) (Figure 3a, iii, iv). This indicates that the elasticity of the granular support bath can be tuned to modulate the extent of shape-morphogenesis in the proposed approach. Interestingly, the ring-shaped constructs exhibited minimal shape-morphing across all packing densities (∼1.5-4% change in diameter) (Figure 3a, iii, v). These lower levels of shape-morphing observed in the bioprinted rings are likely due to the geometric constraints of the closed-loop architecture and the mechanical resistance from the support retained within the core of the ring. In contrast, beam constructs likely experience less structural confinement and can contract more freely along their longitudinal axis.

**Figure 3.**
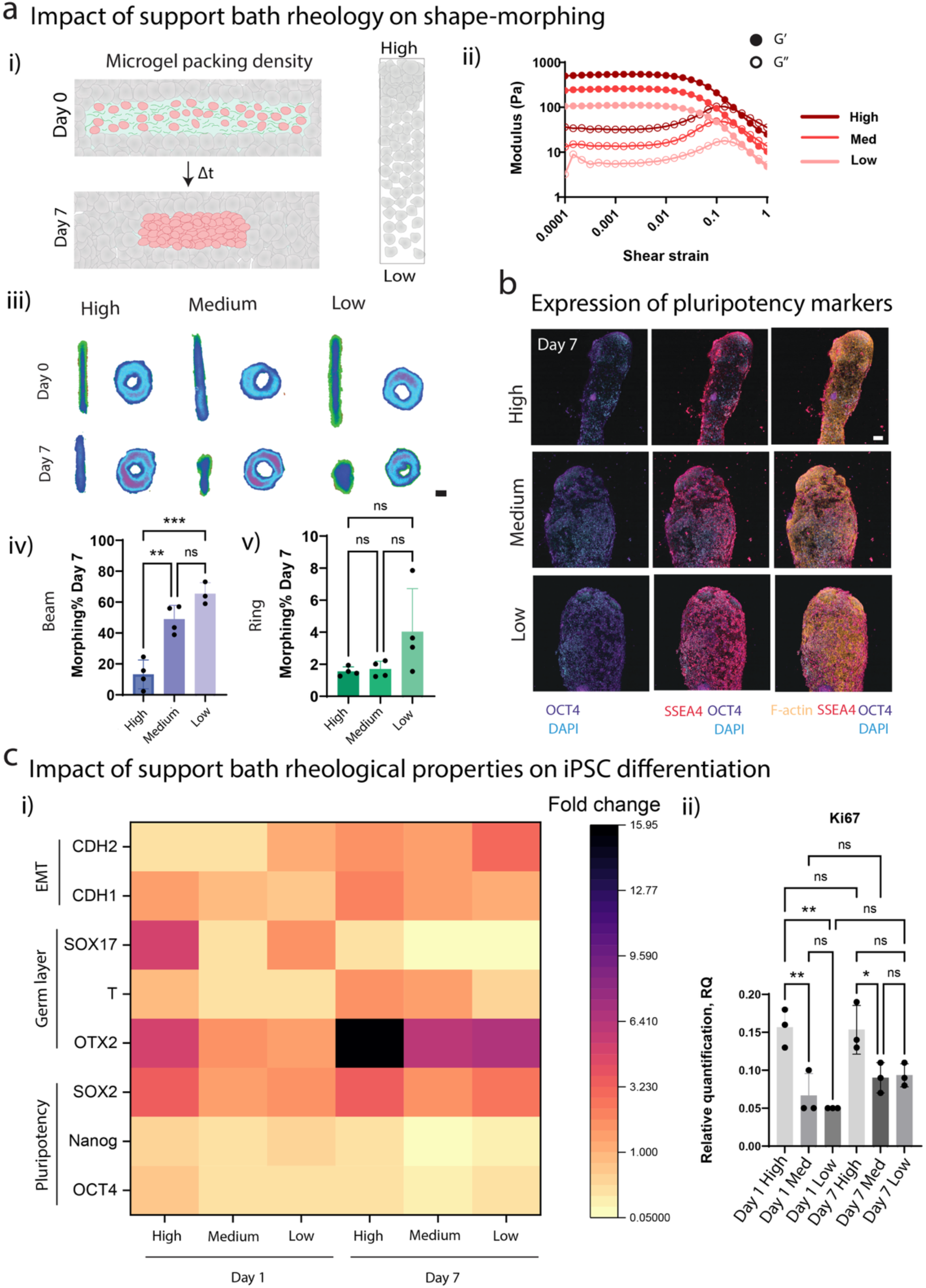
Influence of support bath packing density on tissue shape-morphing: **a)** i) Schematic representation of tissue shrinkage in the granular support bath varying the microgel packing density from high (100%) to low (60%). ii) Amplitude sweep indicated the transition of elastic to fluid-like behaviour of all formulations, modulating the microgel packing density. iii) Brightfield images (scale bar 1 mm) and quantitative analysis showed the influence of varying packing density on shape-morphing in seven-day culture in iv) beam construct and v) ring geometry (n = 3/4 biological replicates, one-way ANOVA with Tukey’s multiple comparison test, where ** denotes p < 0.01 and *** denotes p < 0.001). **b)** Immunofluorescence staining indicated the expression of pluripotency markers (OCT4, SSEA4) in support bath with varying packing densities (scale 100 µm) on day 7. **c)** i) Heatmap demonstrating pluripotency, germ layer, EMT, and proliferation-related gene expression analysis of bioprinted iPSC constructs in high, medium, and low packing densities, with GAPDH as the housekeeping gene and iPSCs cultured in 2D conditions as the control sample (n=3 biological replicates). One-way ANOVA with Tukey’s multiple comparison test was performed for individual genes (Supplementary Figure 3). ii) The plot indicates the fold-change in the proliferation marker Ki67 (relative to iPSCs cultured in 2D conditions, n = 3 biological replicates). One-way ANOVA with Tukey’s multiple comparison test was performed for individual genes (Supplementary Figure 3), and the heatmap is a consolidated version of this data.

### 2.3 Impact of support bath mechanics on iPSC fate in shape-morphing constructs

Next, we examined how the support bath mechanics influenced iPSC pluripotency and differentiation within the shape-morphing constructs. Immunofluorescence staining confirmed the expression of pluripotency markers (OCT4 and SSEA4) up to day 7 (Figure 3b). To further assess iPSC fate, temporal gene expression analysis was performed on days 1 and 7 post-printing using qRT-PCR. The analysis included pluripotency-associated genes (OCT4, Nanog, and SOX2), germ layer-specific genes (OTX2 for ectoderm, Brachyury (T) for mesoderm, and SOX17 for endoderm) and the proliferation marker (Ki67). All results were normalised to those of iPSCs cultured in 2D conditions. The expression of epithelial-to-mesenchymal transition (EMT)-related genes was also evaluated. E-cadherin (CDH1) is essential for iPSC self-renewal, and its expression is associated with the undifferentiated iPSC state ^25^. Conversely, N-cadherin (CDH2) expression is associated with lineage differentiation ^25–27^. The expression of OCT4 and Nanog decreased over 7 days of culture in the 60% packing density support (21% and 70% reduction, respectively) (Figure 3c, i; Supplementary Figure 3a and b). OCT4 and Nanog expression also decreased in the stiffer 100% packing density support (50% and 54% reduction, respectively) (Figure 3c, i; Supplementary Figure 3a, b). SOX2 expression was stable across all formulations. At day 7, the expression of the three pluripotency markers did not differ across the different stiffness supports (Figure 3c, i; Supplementary Figure 3a, b). Notably, the expression of OTX2 (ectoderm marker) increased over 7 days across all stiffness formulations, with the highest expression (fold-change ∼16) observed in the stiffest bath (Figure 3c, i; Supplementary Figure 3a, b). This suggests that the increased physical constraint provided by a stiffer support bath can impact iPSC differentiation within the constructs during morphing. Ki67 expression also increased in the stiffer support bath, indicating enhanced proliferation (Figure 3c, ii). Slight increases in CDH1 expression (E-cadherin, associated with an undifferentiated state) were also observed in the stiffer baths (Figure 3c, i; Supplementary Figure 3a, b). Finally, no significant changes in SOX17 (endoderm) or T-Brachyury (mesoderm) expression were observed between the conditions on day 7 (Figure 3c, i; Supplementary Figure 3a, b).

Next, we investigated whether removing the support bath on day 3 of culture affected iPSC differentiation. As our goal was to subsequently induce cardiac differentiation by modulating the WNT pathway, we sought a bath formulation that maintained pluripotency while minimising spontaneous differentiation. The softer support bath formulation (60% packing density) was selected for these studies, as it had previously resulted in reduced expression of iPSC differentiation markers while enhancing shape-morphing. The rationale for removing the support bath in the later stages of shape-morphing was twofold. First, removal simplifies the delivery of biochemical cues for differentiation. Second, as softer baths imposed less physical resistance to shape-morphing and were associated with reduced spontaneous differentiation, removing the support at later stages was expected to further reduce differentiation and preserve pluripotency. Temporal gene expression analysis was performed on shape-morphing constructs on days 1, 3, and 7. Notably, no significant changes in pluripotency marker expression were observed over 7 days (Figure 4a, iii, Supplementary Figure 4). Additionally, immunofluorescence staining for OCT4 and SOX2 was observed on day 7 (Figure 4a, ii, Supplementary Figure 4b). The expression of germ layer markers also remained unchanged during the culture period (Figure 4a, iii, Supplementary Figure 4b). These results suggest that the iPSC population remained in a pluripotent state when cultured in softer support hydrogels during the early stages of the shape-morphing process. It should be noted that E-cadherin staining (associated with undifferentiated cells) appeared to decrease over 7 days of culture (Figure 4a, ii); however, no significant changes in CDH1 or CDH2 expression were observed by day 7 (Figure 4a, iii, Supplementary Figure 4b). Finally, between days 3 and 7, Ki67 expression increased by 82%, suggesting active iPSC proliferation within the constructs (Figure 4a, iv).

**Figure 4.**
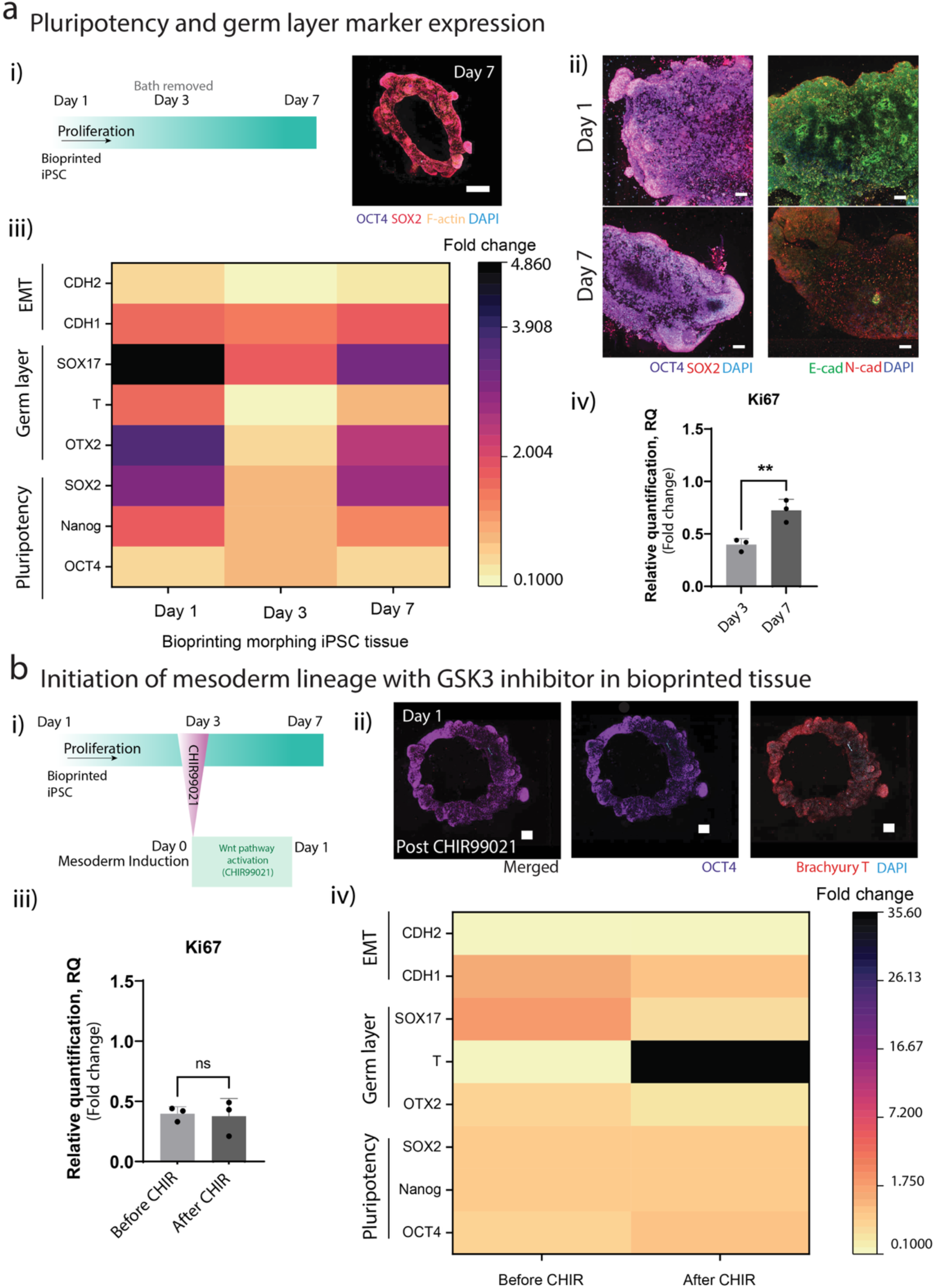
Expression of pluripotency, germ layer, and EMT-associated genes within the bioprinted iPSCs morphing tissue in culture and upon mesodermal induction: **a)** i) Schematic representation of the time points day 1, 3, and 7 to evaluate temporal change in expression of pluripotency (OCT4, Nanog, and SOX2), germ layer (OTX2, T, SOX17), and epithelial-to-mesenchymal transition (CDH1 and CDH2) via qRT-PCR. (Right) Confocal image of a bioprinted ring on day 7 (scale 500 µm). ii) Immunofluorescence images of bioprinted tissue highlighted the expression of pluripotency markers (OCT4 and SOX2) and EMT markers (E-cadherin and N-cadherin) on days 1 and 7 (scale 100 µm). iii) The heat map demonstrates the temporal change in expression during the culture period with GAPDH as the housekeeping gene and iPSCs cultured in 2D conditions as the control sample (n=6 biological replicates). iv) Quantitative analysis of the proliferation marker Ki67 as fold-change value on days 3 and 7 relative to day 1 bioprinted tissue (n = 3 biological replicates, unpaired t-test, where ** denotes p< 0.01). **b)** i) Schematic flow representing the time points selected to evaluate pluripotency, germ layer, and epithelial-mesenchymal transition (EMT) markers before and after WNT activation using CHIR99021. ii) Confocal images of the bioprinted ring upon mesoderm induction on differentiation day 1 showed immunostaining for OCT4 and the mesoderm marker Brachyury (T) (scale 200 µm). iii) The plot indicates the fold-change in the proliferative marker Ki67 (relative to day 1 samples) before and after mesoderm induction on differentiation day 0 (or day 3 post-bioprinting) and day 1 (post-CHIR99021) relative to day 1 post-bioprinted iPSCs (n = 3 biological replicates, unpaired t-test, where ns denotes not significant). iv) The expression of genes associated with pluripotency, germ layer, and EMT was evaluated and represented as a heat map, with GAPDH as the housekeeping gene and 2D iPSCs as the control sample (n=6 biological replicates). Unpaired t-test and one-way ANOVA with Tukey’s multiple comparison test were performed for individual genes (Supplementary Figures 4 and 5). Heatmaps are a consolidated version of this data.

Collectively, these results demonstrate that the stiffness of the support bath can modulate the fate of iPSCs within shape-morphing constructs. The stiffest bath, which restricted morphing, induced a significant upregulation of the ectodermal marker OTX2. Our previous study demonstrated that stiffer granular supports increase the stress and strain within bioprinted shape-morphing constructs ^5^. Mechanical cues have been shown to impact iPSC differentiation ^28,29^, which likely contributed to the elevated expression of OTX2 in stiffer support baths. In contrast, softer support hydrogels supported greater morphing while simultaneously suppressing spontaneous lineage commitment. Furthermore, the removal of the softer support bath after three days resulted in the sustained expression of pluripotency markers without activating differentiation. Overall, these results indicate that the embedded bioprinting of iPSCs within softer granular support baths enables the fabrication of cell-dense, pluripotent 3D constructs with defined architectures. The maintenance of pluripotency throughout the culture period highlights the potential for subsequent directed mesodermal and cardiac lineage differentiation through the biochemical modulation of the WNT pathway during shape-morphogenesis. Accordingly, the next phase of the study focused on developing mesodermal and cardiac differentiation protocols for bioprinted shape-morphing constructs.

### 2.4 Guided *in situ* mesodermal and cardiac lineage differentiation within bioprinted constructs

Building on the findings that pluripotency was preserved during shape-morphing for at least 3 days in softer support bath formulations, we next explored *in situ* mesodermal and cardiac differentiation by modulating WNT signalling. On day 3 post-bioprinting, WNT signalling was activated using the GSK3 inhibitor CHIR99021 to induce mesodermal specification (Figure 4b). To evaluate the effects of WNT activation, qRT-PCR was performed before and after CHIR99021 addition (Figure 4b, iii and iv; Supplementary Figure 5). A significant upregulation (∼35-fold-change) of brachyury (T), a T-box transcription factor and mesodermal marker, was observed (Figure 4b, iv; Supplementary Figure 5). The expression levels of OTX2 (ectoderm) and SOX17 (endoderm) remained unchanged, indicating that differentiation was restricted to the mesodermal lineage only. The expression of E-cadherin, N-cadherin, and Ki67 remained unchanged in response to WNT activation (Figure 4b, iii and iv; Supplementary Figure 5). These results demonstrate the successful induction of mesodermal differentiation *in situ* within shape-morphing tissues, highlighting the potential of using targeted signalling modulation to recapitulate developmental-like iPSC lineage specification in bioprinted constructs.

Next, *in situ* cardiac differentiation was initiated by temporally modulating WNT signalling using CHIR99021 for activation, followed by IWP-4 for inhibition (Figure 5a, i). The resulting tissues were then cultured for up to 21 days, with spontaneous contractions beginning on approximately day 8-12, confirming successful cardiac differentiation (Figure 5b). The bioprinted constructs also underwent shape-morphing during the *in situ* differentiation process, with continued shrinkage of the tubes observed up to day 21 (Figure 5a, ii, iii). To quantify these contractions, the contraction amplitude (an image-based estimation of contraction force) and peak-to-peak time (interval between two contraction peaks) were analysed using the open-source musclemotion software (Figure 5b, i, ii, iii). The constructs exhibited a higher contraction amplitude on day 12, with significant reductions on day 21 (Figure 5b, i). The peak-to-peak time remained constant between days 12 and 21, corresponding to an average beating frequency of ∼0.8 Hz (Figure 5b, ii). This reduction in contraction force over time is potentially due to changes in ECM stiffness during culture. Successful iPSC-CM differentiation was confirmed by positive staining for cardiac troponin T (cTnT) (Figure 6b; Supplementary Figure 6a). Stronger cTnT staining was observed in the peripheral regions of the constructs (Figure 6b), which may be due to the higher concentrations of WNT activators/inhibitors in these regions during differentiation. Additionally, we observed new tissue clusters emerging along the edges of the bioprinted constructs after WNT inhibition, which stained positively for cTnT by the end of the culture period (Figure 6b; Supplementary Figure 6a). The concentration of CHIR99021 plays a key role in iPSC-CM differentiation protocols and must be optimised for each iPSC line. We observed successful iPSC-CM differentiation using both 6 and 7 µM CHIR99021 in our *in situ* protocol.

**Figure 5.**
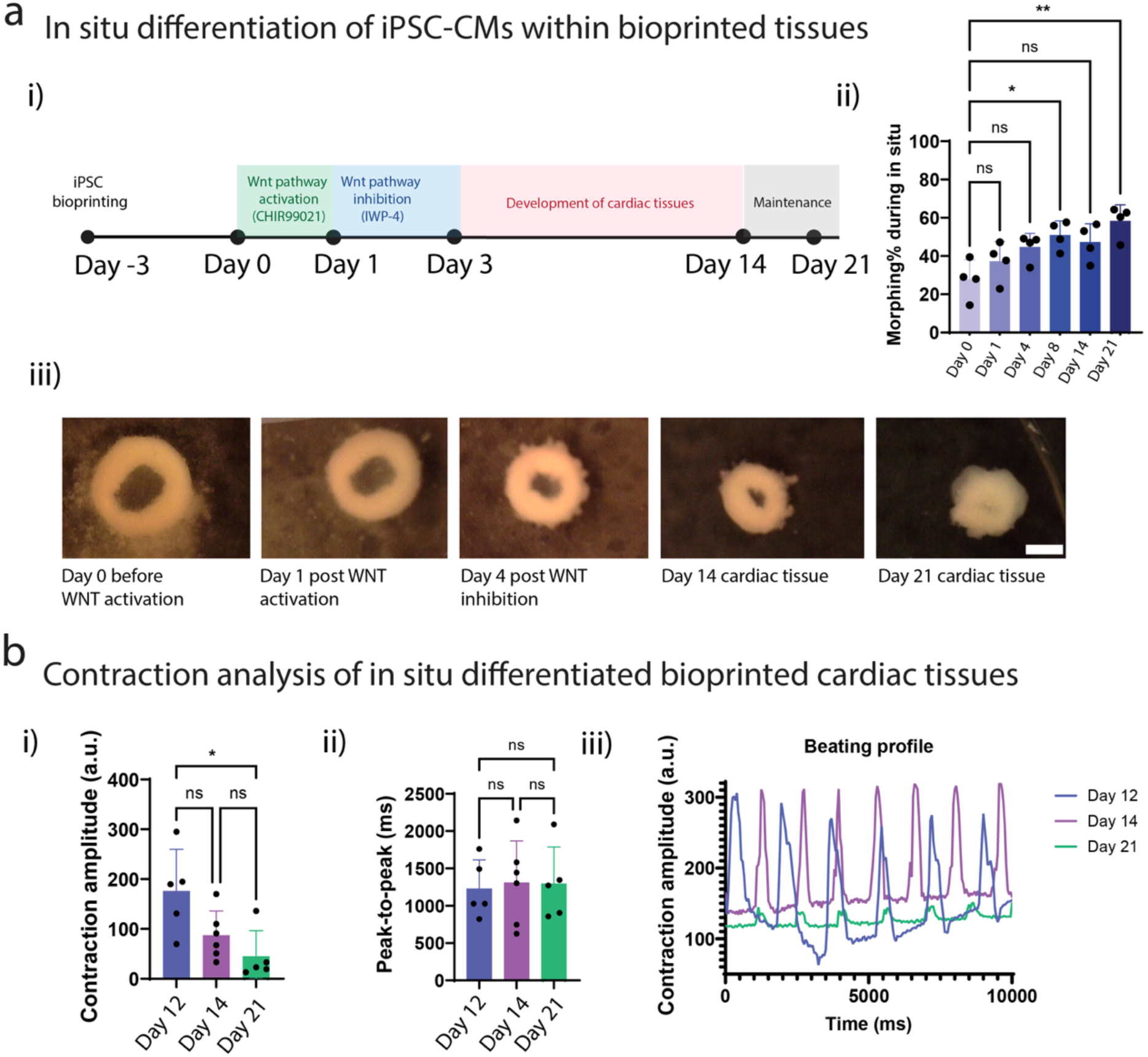
*In situ* differentiation of bioprinted pluripotent shape-morphing constructs into cardiac tissue: **a)** i) Schematic overview of the modified GiWi protocol used to induce mesoderm and cardiac differentiation within bioprinted shape-morphing tissues. ii) Quantitative analysis of shape-morphing during differentiation (n = 3 biological replicates, one-way ANOVA with Tukey’s multiple comparison test, where ns denotes not significant, and * denotes p < 0.05, ** denotes p < 0.01). iii) Brightfield images of bioprinted iPSC tissues before WNT activation and during *in situ* cardiac differentiation (scale 1 mm). **b)** i) Contraction amplitude, ii) peak-to-peak time, and iii) contraction profile of *in situ* differentiated bioprinted cardiac tissue on days 12, 14, and 21 (n = 5/6 biological replicates, one-way ANOVA with Tukey’s multiple comparison test, where ns denotes not significant, and * denotes p < 0.05).

**Figure 6.**
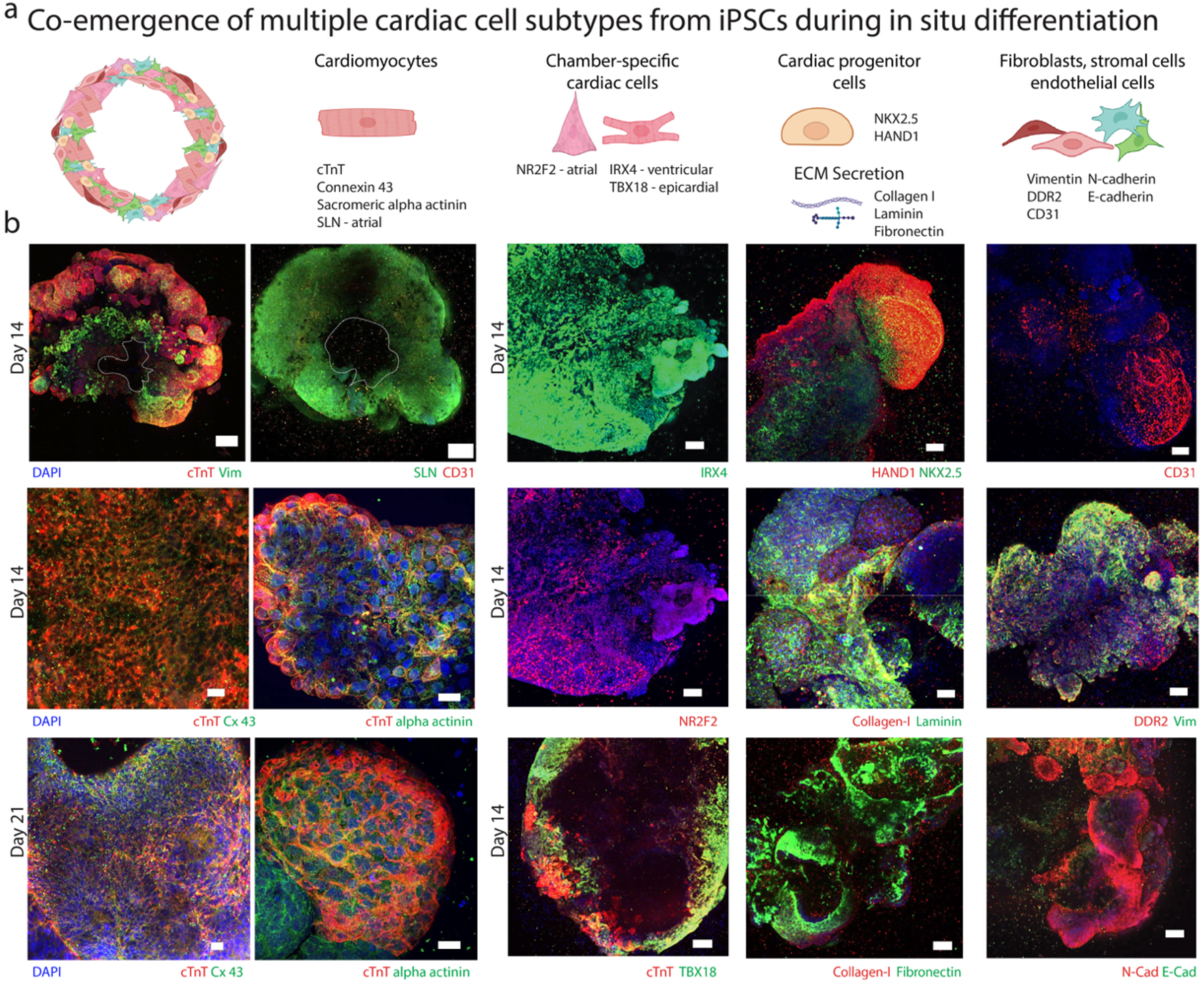
Co-emergence of cardiomyocytes and multiple cardiac cell subtypes from iPSCs during *in situ* differentiation: **a)** i) Schematic illustration (top) and **b)** immunofluorescence images (below) of *in situ* differentiated bioprinted cardiac tissues confirming the emergence of cardiomyocytes and cardiac subtypes using a panel of antibodies on day 14 and day 21. The immunofluorescence analysis includes markers for cardiomyocytes (cTnT, connexin 43, sarcomeric alpha actinin), ventricular cardiomyocytes (IRX4), atrial cardiomyocytes (NR2F2, SLN), epicardial cells (TBX18), cardiac progenitors (HAND1, NKX2.5), fibroblasts (DDR2, vimentin), endothelial cells (CD31), EMT transition markers (E-cadherin, N-cadherin) and ECM secretion (collagen-I, fibronectin, laminin). The lumens of bioprinted ring constructs are indicated using a white dashed mark. (Scales are as follows: 200 µm for the images of the complete ring in b top left, 50 µm for collagen-laminin images, 20 µm for Cx43 and alpha actinin images, and 100 µm for all other images)

Next, we evaluated the presence of cardiomyocyte maturation markers, connexin 43 (Cx43) and sarcomeric alpha actinin. Increased Cx43 staining between days 14 and 21 indicated gap junction formation (Figure 6b). Notably, the iPSC-CMs displayed extensive cell-cell contact throughout the constructs, which is critical for cardiac tissue maturation and function. Sarcomeric alpha actinin staining was observed in regions that stained positively for cTnT (Figure 6b). We also stained for HAND1, a transcription factor marker for cardiac progenitor formation, and NKX2.5, an early marker for cardiac differentiation expressed in linear heart tubes. Positive staining for both HAND1 and NKX2.5 at day 14 confirmed that our *in situ* differentiation protocol initiated embryonic-like differentiation cascades within our bioprinted tissues (Figure 6b). Next, we evaluated whether *in situ* differentiation led to the formation of specific cardiac subtypes. Robust staining for IRX4, a transcription factor marker of ventricular cardiomyocytes, was observed (Figure 6b). We also observed staining for NR2F2 (Figure 6b), a transcription factor marker for atrial cardiomyocytes, although the staining was less intense compared to IRX4, indicating that our *in situ* differentiation protocol predominantly generated ventricular-like iPSC-CMs. Despite the higher IRX4 expression relative to NR2F2, we observed positive staining for sarcolipin (SLN) (Figure 6b), a marker usually associated with atrial cardiomyocytes ^30,31^. These results suggest that a mixed population of ventricular and atrial iPSC-CMs, predominantly of the ventricular phenotype, emerged during *in situ* differentiation. However, staining and functional analyses at later time points would be required to definitively confirm atrial differentiation. Finally, we also observed positive staining for TBX18 (Figure 6b), a marker of epicardial cells ^32,33^.

Next, we evaluated the potential co-emergence of fibroblasts and endothelial cells with cardiomyocytes during *in situ* differentiation. Positive staining for DDR2 (Discoidin Domain Receptor 2), a well-recognised marker for cardiac fibroblasts, together with vimentin (VIM), confirmed the co-emergence of fibroblasts within the shape-morphing constructs (Figure 6b). We also observed increased expression of N-cadherin relative to E-cadherin, a characteristic feature of the cadherin switch associated with cell differentiation during heart development ^34,35^. Fibroblasts were interspersed with cardiomyocytes throughout the tissue, although higher staining was observed in the central regions of the tubes (Figure 6b, Supplementary Figure 6a). This co-emerging fibroblast population likely contributed to the observed shape-morphing during *in situ* differentiation of the cells. For example, our previous work demonstrated that fibroblast-generated contraction forces can drive shape-morphing in bioprinted heart tissues containing co-cultures of pre-differentiated iPSC-CMs and cardiac fibroblasts ^5^. Interestingly, we also observed positive staining for CD31 at day 14, indicating that endothelial cells co-differentiated with cardiomyocytes (Figure 6b). These results demonstrate that our *in situ* differentiation protocol promoted the co-emergence of cardiomyocytes, fibroblasts, and endothelial cells, resembling the multicellular composition of developing cardiac tissue. This cellular heterogeneity is likely influenced by the 3D microenvironment, which will induce spatial gradients in small-molecule concentrations during differentiation. Additionally, the dynamic shape-morphing process likely modulates local ECM mechanics, thereby altering cellular interactions with the surrounding matrix during in situ differentiation. For example, positive staining for collagen-I, fibronectin, and laminin demonstrated nascent ECM deposition by the embedded cells during differentiation (Figure 6b). ECM proteins such as fibronectin and laminin have been shown to influence iPSC-CM differentiation ^36^.

To assess transcriptional changes associated with cardiac differentiation and maturation, qRT-PCR was performed on days 14 and 21 (Figure 7a, Supplementary Table 1). Genes associated with cardiac muscle development (*MYH6, MYH7, and MYL7*), cardiac muscle contraction (*TNNI3, TNNT2, and TNNI1*), ion channels (*ATP2A2, SCN5A, CACNA1C, and KCNJ2*), gap junctions (*GJA5*), cardiac transcription factors (*NKX2.5 and GATA4*), and stromal cell identity (*DDR2, ACTA2, POSTN, and VIM*) were significantly upregulated at days 14 and 21 relative to pre *in situ* differentiation (Figure 7a). All significant changes in gene expression and associated fold-change values are listed in Supplementary Table 1. The upregulation of cardiac transcription factors *NKX2.5* and *GATA4* confirmed early cardiac lineage commitment within the tubes, while increases in TNNI1 (troponin 1, skeletal, slow isoform), *TNNT2* (cardiac troponin T type 2), and *TNNI3* (troponin I cardiac isoform) further validated cardiomyocyte differentiation. The lower fold-change increase of *TNNI3* relative to *TNNI1* indicates that the *in situ* differentiated constructs mimic an early cardiac developmental stage. The upregulation of *MYH7* (beta-myosin heavy chain), *MYL7* (myosin light chain atrial isoform), and *MYH6* (alpha-myosin heavy chain) at day 14 indicated that the nascent cardiac muscle was maturing, and further significant increases in *MYL2* and *MYH7* expression at day 21 suggested continued cardiac tissue maturation within the bioprinted constructs as the culture period was extended. Genes encoding key cardiac ion channels, including *KCNJ2* (potassium ion transport), *CACNA1C* (L type voltage-gated calcium), *SCN5A* (sodium influx), and *ATP2A2* (calcium regulation), increased 2-3-fold between days 14 and 21, indicating enhanced electrophysiological maturity with longer culture periods. Interestingly, despite the use of an adapted ventricular differentiation protocol, *GJA5* (connexin 40, atrial gap junction) expression doubled by day 21, which was consistent with positive staining for SLN. Fibroblast and stromal cell-associated genes (*POSTN, ACTA2, DDR2, and VIM*) were also upregulated by day 14, further supporting the co-emergence of cardiac fibroblasts alongside cardiomyocytes. By day 12, a subset of constructs exhibited spontaneous contractions (categorised as beating samples), whereas others remained non-beating by day 21. Comparative gene expression analysis between these groups at day 21 demonstrated that the beating samples expressed higher levels of cardiac transcription factors (*NKX2.5* and *GATA4*), muscle contraction/development genes (*MYL7, MYH6, MYH7, TNNI1, TNNT2, and TNNI3*), and ion channels (*ATP2A2, SCN5A*) (Figure 7b, Supplementary Table 2).

**Figure 7.**
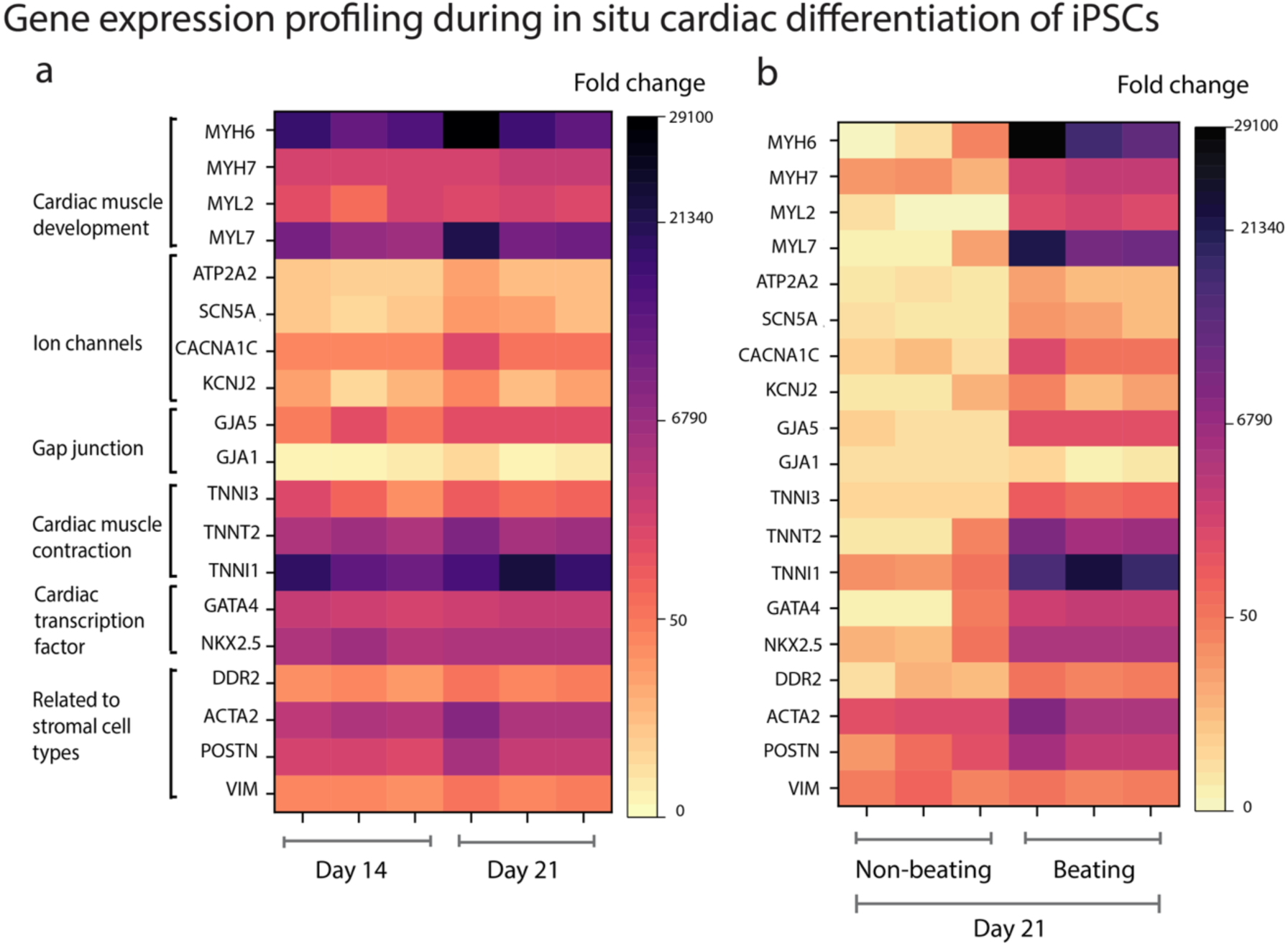
Transcriptomic profile of *in situ* differentiated bioprinted cardiac tissue: **a)** Representation of quantitative fold-change as a heatmap of *in situ* differentiated bioprinted cardiac tissue at days 14 and 21, and **b)** at day 21 categorised based on their contractile properties, where non-beating refers to the lack of spontaneous contractile forces observed post-*in situ* differentiation. GAPDH was the housekeeping gene, and undifferentiated (before WNT activation) bioprinted iPSCs were the control samples (n = 3 biological replicates). A list of significantly upregulated genes is presented in tabular form in Supplementary Tables S1 and S2.

These findings collectively demonstrate that bioprinted pluripotent tissues can be guided through mesodermal and cardiac differentiation *in situ* via WNT pathway modulation. Unlike traditional cardiac bioprinting strategies that employ 2D pre-differentiated iPSC-CMs, our method induces cardiac differentiation post-printing as the tissue undergoes shape-morphing. By aligning differentiation and morphogenesis, this strategy more closely emulates embryonic heart development, in which cardiac progenitors and cardiomyocytes emerge within the cell-dense, matrix-rich, 3D microenvironment of the looping heart tube. As noted in the introduction, one prior study has reported *in situ* cardiac differentiation of iPSCs within bioprinted constructs ^9^. However, in this study, the authors reported non-uniform cardiac differentiation that was restricted to the peripheral regions of the construct, with the core areas remaining largely acellular. This spatial heterogeneity may have arisen from the use of a stiffer, covalently crosslinked collagen/GelMA-based bioink, which could limit iPSC survival in the core regions. Although the final dimensions of our bioprinted rings were small (∼1.4 mm diameter by day 21), we observed uniform cell densities and relatively homogenous cardiomyocyte differentiation throughout our constructs, as evidenced by cTnT, Cx43, and alpha actinin staining (Figure 6b). This uniformity was likely facilitated by the high cell density used for bioprinting (150 million cells mL^−1^) and cell compaction during shape-morphing. Additionally, our embedded bioprinting approach enabled the use of a soft Matrigel bioink that supported cell proliferation and self-organisation. It should also be noted that the mechanical forces acting on iPSCs during shape-morphing may have also influenced cardiac differentiation trajectories, analogous to shape-transformations of the heart tube that produce mechanical stresses in the tissue that coincide with mesoderm-to-cardiac transitions ^16,17^.

Our *in situ* differentiation approach resulted in the co-emergence of cardiomyocytes, cardiac fibroblasts, and endothelial cells within a single bioprinted construct, which has not been reported previously. While co-culture strategies have been shown to enhance iPSC-CM differentiation ^5,37,38^, they typically involve mixing pre-differentiated cell populations. In contrast, our *in situ* differentiation approach enables the co-emergence of multiple cardiac cell types from a common progenitor pool within a dynamic, shape-morphing microenvironment. The resulting cellular crosstalk, which occurs when cells are in a more developmentally plastic state, likely affects cardiac differentiation and maturation trajectories. Positive staining for early cardiac transcription factors, such as NKX2.5 and HAND1, confirmed the progressive transition from cardiac progenitor to cardiomyocyte phenotypes within our bioprinted constructs. Crosstalk among cells at varying differentiation stages may further influence the maturation trajectories. In contrast, in traditional approaches using pre-differentiated iPSC-CMs, these developmental transitions occur in 2D culture conditions outside the bioprinted construct. Consequently, such constructs may lack instructive microenvironmental cues that promote structural and functional maturation. Collectively, our results highlight the potential of combining shape-morphing and in situ differentiation to generate nascent cardiac tissues that can mature during culture via endogenous developmental programmes.

## 3. Conclusion

This study presents a novel developmentally inspired strategy for bioprinting human heart tissue by directing the *in situ* cardiac differentiation of iPSCs within shape-morphing constructs. The embedded bioprinting of Matrigel bioinks containing high-density suspensions of iPSCs into granular support baths, followed by post-printing cellular condensation, generated pluripotent, cell-dense tissues capable of undergoing shape-morphing. Support bath mechanics were optimised to maintain pluripotency, with softer formulations minimising spontaneous differentiation while allowing shape-morphing. Temporal modulation of WNT signalling guided mesodermal and cardiac differentiation within the shape-morphing constructs, generating nascent human heart tissues in which cardiomyocytes, fibroblasts, and endothelial cells co-emerged from a common progenitor pool. By aligning morphogenesis with mesoderm and cardiac differentiation, this approach recapitulates key aspects of early heart development, where cardiac lineage specification occurs within the looping heart tube. Overall, this developmentally inspired platform represents a significant departure from conventional 2D pre-differentiation paradigms and offers a promising pathway for bioprinting nascent heart tissues with an endogenous capacity for structural and functional maturation.

## 4. Experimental section

### Human induced pluripotent stem cell (iPSC) culture, maintenance, and expansion

Human iPSCs (A18945, Fisher Scientific) were seeded on Corning^®^ Matrigel^®^ (354277, Fisher Scientific) coated 6-well plates. The Matrigel coating solution was prepared by diluting Matrigel (2%) in Gibco^TM^ DMEM F-12 GlutaMAX^TM^ supplement medium (10565018, Fisher Scientific), and the plates were coated for 1 h at room temperature, followed by incubation at 37 °C for 20 min. 5 µM ROCK Inhibitor Y-27632 (72304, STEMCELL Technologies), was added to the culture medium during thawing to enhance cell survival. iPSCs were detached using Accutase^TM^ (07920 or 07922, STEMCELL Technologies) for 5-7 min in incubation at 37 °C. Detached cells were centrifuged at 300G for 5 min, and the cell pellet was gently dissociated to seed them either on coated well plates or embedded in ink. iPSCs were maintained in a 1:1 ratio of mTeSR^TM^ Plus (100–1130, STEMCELL Technologies) and Gibco Essential 8^TM^ medium (A2858501, Fisher Scientific) for the first day after thawing and passaging, followed by daily changes of Gibco Essential 8^TM^ medium until the cells reached 80–90% confluency. For iPSC bioprinting, cells were used between passages 9-12. ROCK inhibitor (10 µm concentration) was added to the Gibco Essential 8^TM^ medium for 24h post-bioprinting to enhance cell survival. For the first three days, the media was changed daily; after three days, it was changed on alternate days.

### Pre-formed embryoid bodies and bioprinting

iPSCs were passaged once the cells reached 80–90% confluency. The cell suspension (2.5 million cells per well) was reseeded in an AggreWell™ 400 6-well plate (34425, STEMCELL Technologies) after passaging. Before seeding, the plate was coated with an anti-adherence solution (07010, STEMCELL Technologies) for 5 min and centrifuged at 1300G for 5 min. This was followed by rinsing with warm culture medium once and then storing in an incubator with culture medium unless the cells were passaged. After seeding the cells, the plate was centrifuged at 100G for 3 min. For the first day, a 1:1 ratio of mTeSR^TM^ plus medium and Gibco Essential 8^TM^ medium were added (total 4ml each well), followed by Essential 8^TM^ medium the next day. Pre-formed EB were aspirated using a cut pipette after two days and encapsulated in a Matrigel-based bioink for bioprinting. ROCK inhibitor (10 µM) was added for the first 24h, followed by Gibco Essential 8^TM^ medium. For the first three days, the media was changed daily; after three days, it was changed on alternate days.

### In situ differentiation to cardiac tissue

Bioprinted iPSC tissues were maintained in Gibco Essential 8^TM^ medium with daily media changes and gradual agarose removal over three days. On day 3, iPSC-laden tissue (from passages 9 and 11) was directed towards the mesoderm lineage by adding 6-7µM CHIR99021 (72054, STEMCELL Technologies) and removing the agarose support bath, which was considered day 0 of differentiation. This was followed by 1µM CHIR99021 on day 1, 5µM IWP-4 (72554, STEMCELL Technologies) on day 3. These small molecules were added to Gibco RPMI 1640 medium (11875085, Fisher Scientific) supplemented with Gibco B27 minus insulin (2%) (A1895601, Fisher Scientific). This was followed by medium change without any small molecules on day 5 of differentiation. Next, the culture medium was changed to RPMI 1640 supplemented with Gibco B27 supplement (2%) (17504044, Fisher Scientific) and 1% penicillin-streptomycin on day 8 and changed every 2 days until day 14.

### Bioink Preparation

Type I bovine atelocollagen (PureCol^®^ 3 mg mL^−1^ from Advanced Biomatrix 5005) was neutralised according to the manufacturer’s instructions to adjust the pH to ≈7.2. The final concentration after neutralisation was 2.4 mg mL^−1^, and the collagen was then diluted to a final concentration of 0.96 mg mL^−1^ for cell encapsulation and printing (by volume; 40% collagen, 60% media and cells). The bioink was stored in an icebox to prevent gelation before cell encapsulation and bioprinting. Corning® Matrigel® (354277, Fisher Scientific) of dilution factor 260μl (Lot no. 12224005) was used for all studies in this work. According to the manufacturer’s specifications, the typical protein concentration of the Corning® Matrigel® matrix is 8-11 mg ml^−1^. For the preparation of our Matrigel bioink, the final Matrigel protein concentration was diluted to approximately 3.2-4.4 mg mL^−1^ (by volume; 40% Matrigel, 60% media and cells). To prepare the composite collagen and Matrigel bioinks, the final concentrations of collagen and Matrigel were 0.6 mg mL^−1^ and 2-2.75 mg mL^−1^, respectively (by volume; 25% collagen, 25% Matrigel, 50% media and cells).

### Agarose Support Bath Fabrication

Agarose type I, low EEO was purchased from Sigma-Aldrich (A6013). Agarose microparticles were prepared using a shearing technique to form a viscoelastic suspension bath with varied stiffness ^39,40^. Briefly, a 0.5 wt.% agarose solution (1X PBS) was autoclaved at 121 °C, and then the molten solution was allowed to cool at room temperature for 3 h under stirring at 700 rpm. This protocol results in the formation of microparticles via shearing as the agarose solidifies. The agarose microparticle solution was maintained under sterile conditions and stored for up to 3 months at 4–7 °C. Prior to iPSC bioprinting, agarose particles were centrifuged at 900G for 10 min and resuspended in the same volume of E8 medium supplemented with 1% penicillin-streptomycin.

### 3D Bioprinting Setup and Embedded Bioprinting

Bioprinting experiments were performed using an in-house designed bioprinter, Bioframe (Supplementary Figure 1 and b). Bioframe is a desktop core XY 3D printer that supports a high degree of modifications for specific tasks. It features swappable tool heads and print beds (Supplementary Figure 1 d), a large vertical printing range, and runs on the open-source 3D printer firmware, Klipper, allowing for the integration of additional electronic components, such as cameras and light sources. For these experiments, the Bioframe was configured with a deep vessel for printing tall structures and a Puredyne progressive cavity pump extruder (Supplementary Figure 1 c). The Bioframe was placed inside a biosafety hood to maintain sterile conditions (Supplementary Figure 1 e). Models for printing were prepared using computer-aided design (CAD) software, Autodesk Inventor, and G-code for tool path planning of the prints was generated using Prusa Slicer on a custom profile for the Bioframe (Supplementary Figure 1 f). This profile was tuned for use with 27G needles, where the desired extrusion should be of a similar width to the needle and the printer should operate at a speed of 1mm/s. Live adaptation was performed on the prints via the Klipper user interface installed on the Bioframe, by adjusting the extrusion factor to 120% – 150% of the extrusion values prescribed by the slicer in order to achieve optimal print quality. During bioprinting, the material temperature in the extruder was maintained at 6-7 °C using a Puredyne temperature control unit, and post-bioprinting, the constructs were incubated immediately at 37 °C. High glass bottom μ-Dishes (35 mm, 81158, ibidi) were used to culture the bioprinted constructs. These dishes enabled imaging of the tissue within the support bath during the culture period using an inverted microscope. The culture dishes were adapted to contain two compartments separated by a solid agarose divider (2 wt%). Culture medium was added to one compartment, and an agarose suspension bath was added to the second compartment ^39^. The agarose divider allowed the diffusion of the culture medium into the bioprinted constructs suspended in the support bath. The agarose suspension bath supported the bioprinted tissue for the first three days of culture and was then removed for longer culture as well as for *in situ* differentiation (except for the bath stiffness study, where the bath was not removed until day seven). The media was changed daily for the first three days and then on alternate days.

### Rheological Characterisation

The rheological properties of the varying bath stiffness were characterised using rotational shear rheometry (25 mm diameter, gap 1 mm; Anton Paar MCR 302) at room temperature. Storage and loss modulus were measured separately as a function of shear strain (0.0001^−1^–1^−1^).

### Microscopy Analysis of Bioprinted Constructs

A brightfield microscope (Dinocapture^TM^ 2.0) was used to visualise live samples during the culture period. Images of the bioprinted constructs were captured at different time points, and FIJI software was used to measure the shape change relative to day 0. Inverted confocal microscopy was used to image bioprinted samples containing fluorescein isothiocyanate-conjugated collagen and cell dye. Invitrogen^TM^ CellTracker^TM^ dyes (C34552:15 µM, Fisher Scientific) were used to track shape changes. Bioprinted tissues were visualised live on the day of bioprinting (day 0), and fluorescence images were obtained using multi-channel confocal microscopy (Andor benchtop BC43).

### Live/Dead Staining

Live/dead staining was performed by treating samples with 2 µM Calcein AM (11564257, Fisher Scientific) and 2 µM ethidium homodimer (10184382, Fisher Scientific) in 1X PBS, followed by 30-minute incubation at 37 °C. The samples were imaged using an Andor benchtop BC43 confocal microscope. This study quantified the percentage of live cell area (Viability = Live cell area/ (live + dead cell area) as it was difficult to count individual live cells from these images due to the high levels of cell-cell contact.

### Phalloidin TRITC Staining

To visualise the actin cytoskeleton, fixed bioprinted constructs were stained with 0.05 mg mL^−1^ phalloidin tetramethylrhodamine B isothiocyanate (P1951, Merck) for 2 h. The constructs were then washed twice with 1X PBS, followed by staining with Fluoroshield^TM^ DAPI for 20 min at room temperature, and the samples were visualised using confocal microscopy.

### Immunofluorescent Staining

Samples for immunofluorescent staining were fixed with 10% neutral buffered formalin solution overnight at 4 °C and stored in DPBS the following day until staining. Samples were then permeabilised with Triton-X 100 (0.2%) in PBS and 2% BSA for 45 min and then blocked using 2% BSA in PBS for 1 h. Primary antibody (1:200 dilution in blocking buffer) staining was then carried out overnight at 4 °C, followed by three washes with PBS and then adding the secondary antibody for 2 h (1:200 dilution in blocking buffer). Samples were washed thrice before counterstaining with Fluoroshield^TM^ DAPI (F6057, Merck) for 20 min. Imaging was performed using an Andor benchtop BC43 confocal microscope. All primary and secondary antibodies are listed (Supplementary information S3).

### RNA Isolation and qPCR

The PureLink RNA Mini Kit (10307963, Invitrogen^TM^) was used to isolate RNA from 2D iPSC, bioprinted iPSC, and iPSC-CM samples. cDNA preparation was carried out using the Invitrogen™ SuperScript™ IV VILO™ Master Mix (15523145) or RT2 First Strand Kit (330404, Qiagen) and processed in a Veriti Thermal Cycler. RT-PCR was performed using the Applied Biosystems™ TaqMan™ Fast Advanced Master Mix for qPCR (11380912) or RT2 SYBR Green ROX qPCR Mastermix (330522, Qiagen) for a custom-designed RT2 PCR Array 96-well plate (330171, Qiagen), maintaining a 10% excess amount for the qPCR run. QuantStudio 5 was used to run the qPCR, and data analysis was performed in Excel using the CT value. Heatmaps were plotted in OriginPro 2023 software based on the fold-change value calculated using the CT value, considering GAPDH as the housekeeping gene for all studies. Further details on the genes can be found in Supplementary Information S3.

### Statistical Analysis

All experimental data were compiled and analysed using Microsoft Excel. All graphs are presented as means with standard deviation, along with sample numbers denoted in each graph and figure legends. The reported sample sizes (n-numbers) in the figure legends denote biologically independent samples. Statistical analyses were performed using GraphPad Prism 10 software. Unpaired t-test, two-way or one-way ANOVA tests were used depending on the number of independent variables within the experiment, and Tukey’s multiple comparison test was used to compare differences between means. P values are described as follows: ns denotes not significant, * p <0.05, ** p < 0.01, *** p < 0.001, **** p < 0.0001.

## Acknowledgements

This publication has emanated from research conducted with the financial support of the Irish Research Council (GOIPG/2022/485), the European Research Council (Grant number 101077900), and the EU Commission Recovery and Resilience Facility under the Research Ireland Future Digital Challenge Grant Number 22/NCF/FD/10991. This publication has also emanated from research supported in part by a grant from Research Ireland and is co-funded under the European Regional Development Fund under Grant number 13/RC/2073_P2. We would like to acknowledge the University of Galway College of Science and Engineering Postgraduate Research Scholarship Scheme for their support. We would like to acknowledge the Genomics and Screening Core Facility for providing access to PCR equipment. We express our gratitude to Vasileios Sergis for his support in the design of the heart CAD file and Akash Garhwal for his assistance with cell maintenance in one of the studies.

## Data availability statement

All data supporting the results of this study will be made available at the time of journal publication.

## Table of Contents Image and Text

A developmentally inspired bioprinting approach enables the fabrication of pluripotent tissues that undergo shape-morphing and in situ cardiac lineage specification. This method employs embedded bioprinting to deposit iPSCs within soft granular hydrogels to create pluripotent tissue constructs that undergo cell-mediated shape morphogenesis. Temporal modulation of WNT signalling directs mesodermal induction and cardiogenesis directly within the morphing constructs, generating nascent human heart tissues in which cardiomyocytes, fibroblasts, and endothelial cells co-emerge from a common progenitor pool.

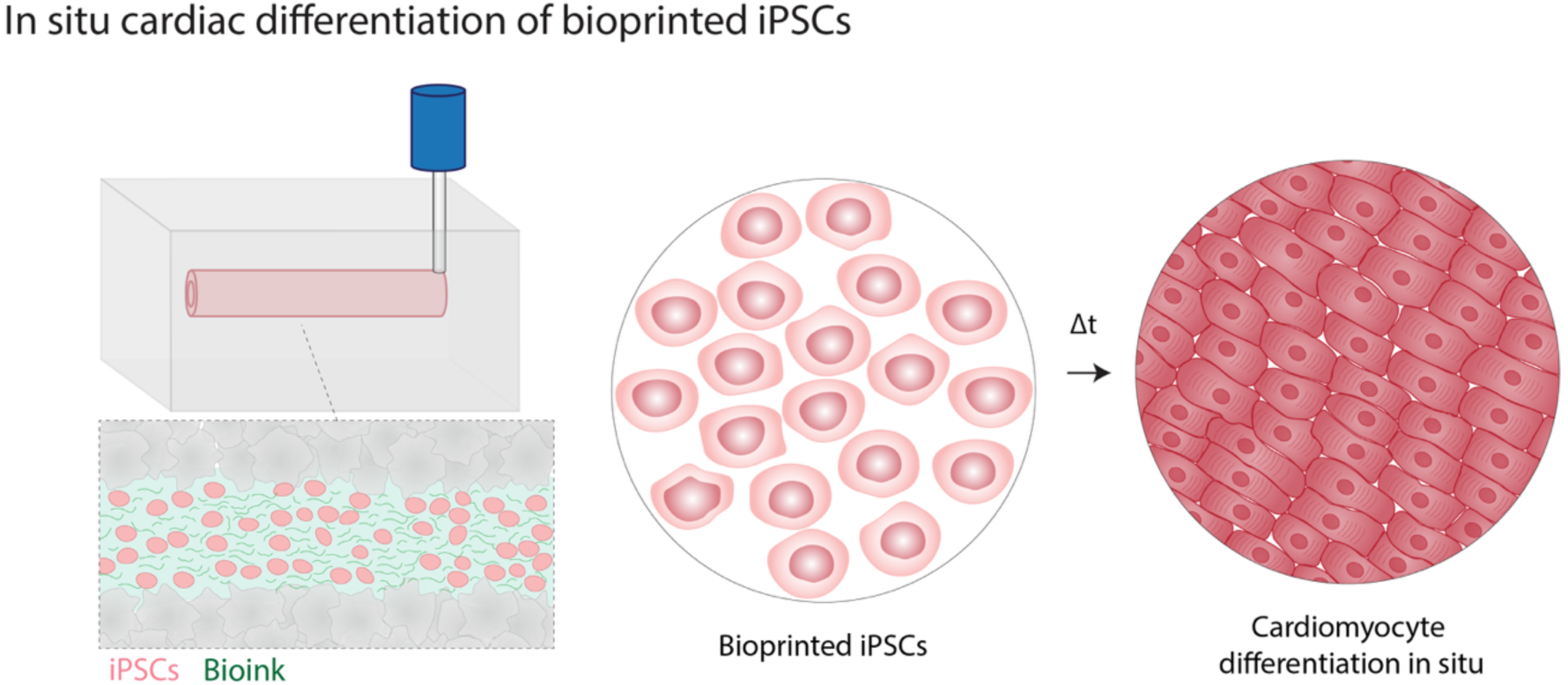

## Supporting Information S1 – Additional Table

**Table S1.1:** Overview of genes that were significantly upregulated at Days 14 and 21 within *in situ* differentiated constructs relative to before *in situ* differentiation. Fold change values represent gene expression values at Day 14/21 relative to before *in situ* differentiation, with GAPDH as the housekeeping gene. At Day 21, the constructs were separated into beating and non-beating groups. Statistical analysis was based on an unpaired t-test, and significant genes were selected based on p< 0.05. All gene expression fold-change values are presented as heatmaps in Figure 7. A statistical comparison between the beating and non-beating samples is presented in Table S1.2.

| **D14**<br>**Beating** | **Fold change** | **D21**<br>**Beating** | **Fold change** | **D21**<br>**Non-Beating** | **Fold change** |
| --- | --- | --- | --- | --- | --- |
| POSTN | 422.85 | VIM | 44.63 | VIM | 50.00 |
| ACTA2 | 3108.47 | POSTN | 2846.21 | POSTN | 61.52 |
| DDR2 | 27.35 | ACTA2 | 5844.57 | ACTA2 | 198.91 |
| NKX2-5 | 4420.90 | DDR2 | 42.99 | DDR2 | 6.32 |
| GATA4 | 775.21 | NKX2-5 | 4314.28 | TNNI1 | 34.64 |
| TNNI1 | 15109.84 | GATA4 | 997.77 | TNNI3 | 4.59 |
| TNNT2 | 4880.00 | TNNI1 | 19284.87 | GJA1 | 3.87 |
| TNNI3 | 125.93 | TNNT2 | 7327.01 | GJA5 | 3.90 |
| GJA5 | 77.89 | TNNI3 | 75.42 | CACNA1C | 5.35 |
| KCNJ2 | 7.87 | GJA5 | 142.92 | ATP2A2 | 2.84 |
| CACNA1C | 35.21 | KCNJ2 | 16.72 | MYH7 | 16.20 |
| SCN5A | 5.49 | CACNA1C | 121.46 |  |  |
| ATP2A2 | 5.94 | SCN5A | 11.37 |  |  |
| MYL7 | 8257.74 | ATP2A2 | 8.91 |  |  |
| MYH7 | 400.26 | MYL7 | 14695.61 |  |  |
| MYH6 | 16185.97 | MYL2 | 285.22 |  |  |
|  |  | MYH7 | 1191.22 |  |  |
|  |  | MYH6 | 20566.46 |  |  |

**Table S1.2:** Overview of genes that were significantly upregulated in ‘beating’ relative to ‘non-beating’ constructs at day 21. As per Table S1.1, fold change values represent gene expression values at Day 21 relative to before *in situ* differentiation, with GAPDH as the housekeeping gene. Statistical analysis was based on an unpaired t-test, and significant genes were selected based on p< 0.05. All gene expression fold-change values are presented as heatmaps in Figure 7.

| Gene | D21 Non-beating (fold change) | D21 Beating (fold change) |
| --- | --- | --- |
| ACTA2 | 198.910 | 5844.570 |
| DDR2 | 6.321 | 42.987 |
| NKX2-5 | 22.511 | 4314.276 |
| GATA4 | 14.568 | 997.767 |
| TNNI1 | 34.637 | 19284.866 |
| TNNT2 | 11.694 | 7327.005 |
| TNNI3 | 4.590 | 75.416 |
| GJA5 | 3.900 | 142.917 |
| SCN5A | 2.942 | 11.372 |
| ATP2A2 | 2.843 | 8.908 |
| MYL7 | 5.987 | 14695.608 |
| MYH7 | 16.198 | 1191.218 |
| MYH6 | 11.731 | 20566.457 |

## Supporting Information S2 – Additional Figures

**Supplementary figure 1.**
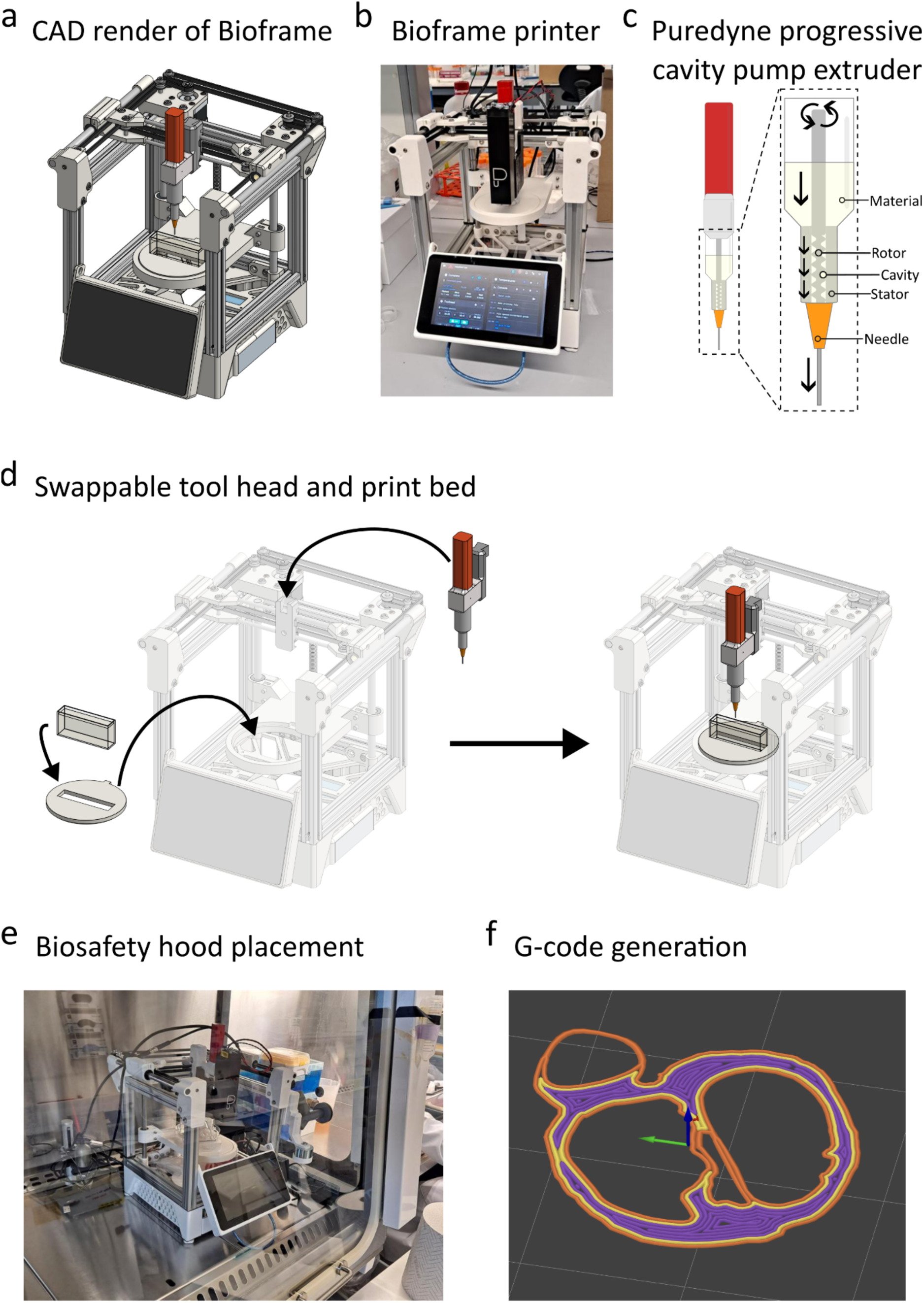
Custom-built bioprinting hardware specifications and operation: **a)** CAD render of Bioframe bioprinter. **b)** Final assembled Bioframe featuring core XY kinematics, temperature-controlled extruder, and Klipper firmware with Mainsail user interface. **c)** Puredyne progressive cavity pump extruder used for bioprinting. **d)** Mechanism for ease of changing tool heads and printbeds on the Bioframe. **e)** Installation of the Bioframe unit in the biosafety hood for sterile printing conditions. **f)** Toolpath rendering from the generated G-code using PrusaSlicer.

**Supplementary figure 2.**
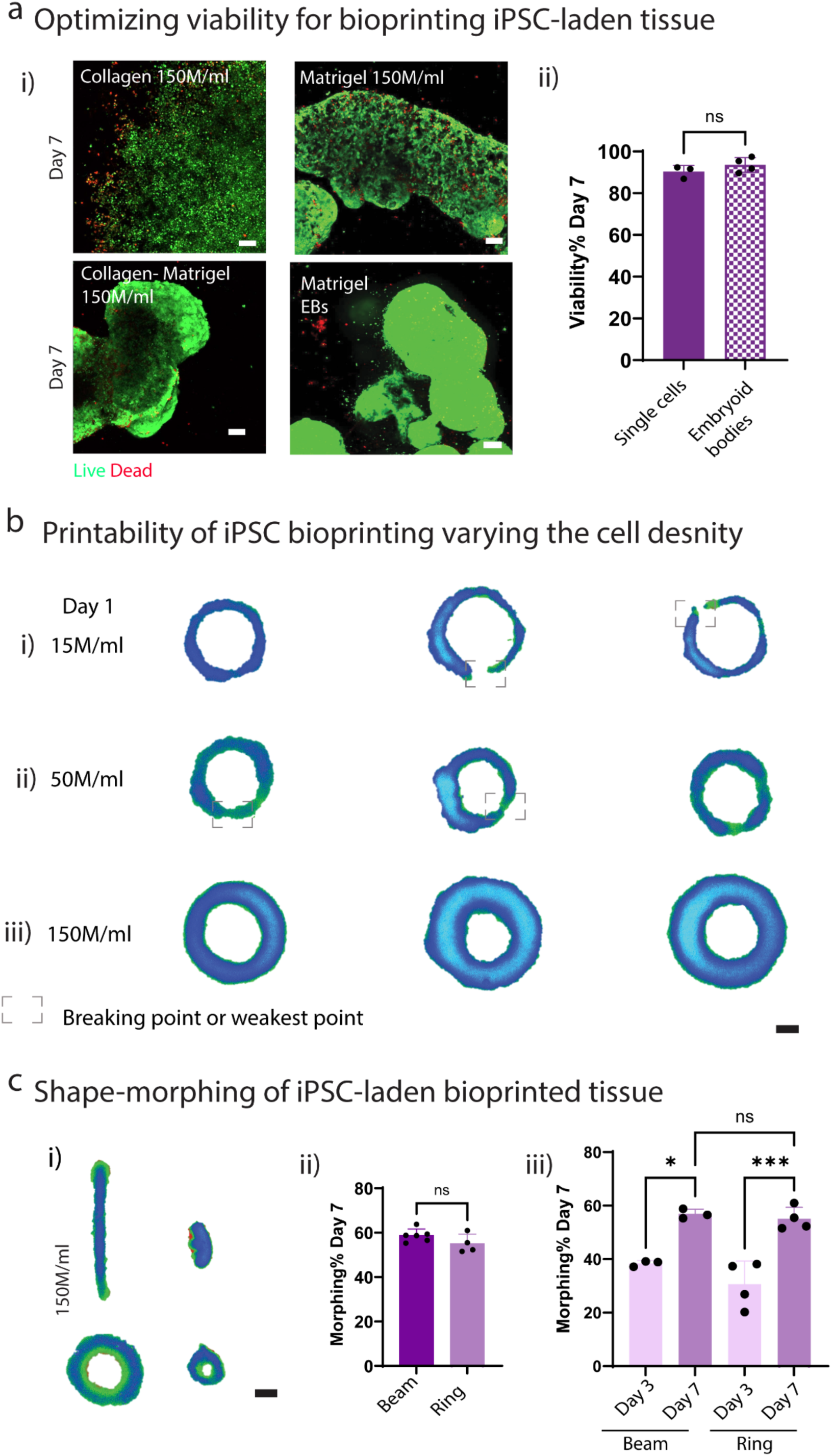
Optimising viability and printability of bioprinted iPSCs: **a)** i) Confocal images of live-dead viability staining for different bioink compositions and cell entities (scale bar 100 µm), ii) Quantitative analysis demonstrating the viability of bioprinted tissue on day 7, varying the cell entity within bioink (single cells vs. embryoid bodies). **b)** Printability of bioprinted constructs was evaluated through brightfield images post 24h bioprinting to identify the weak points or breakage along the fabricated ring for i) 15, ii) 50, and iii) 150 million cells ml-1 (scale bar 1 mm). All biological replicates n=3/4, an unpaired t-test was performed, where ns denotes no significance. **c)** i) Brightfield images highlighted the tissue shape-morphing in culture for beam and ring constructs with 150 million cells mL^−1^ (scale bar 1 mm). ii) Quantitative analysis demonstrated the extent of shape-morphing in beam and ring constructs over seven days of culture. iii) The Beam and Ring constructs demonstrated a significant extent of shape-morphing from day 3 to day 7, which was analysed quantitatively from brightfield images using Fiji software. All biological replicates (n=4-6), one-way ANOVA was performed where ns denotes not significant, * denotes p <0.05, and *** denotes p< 0.001.

**Supplementary figure 3.**
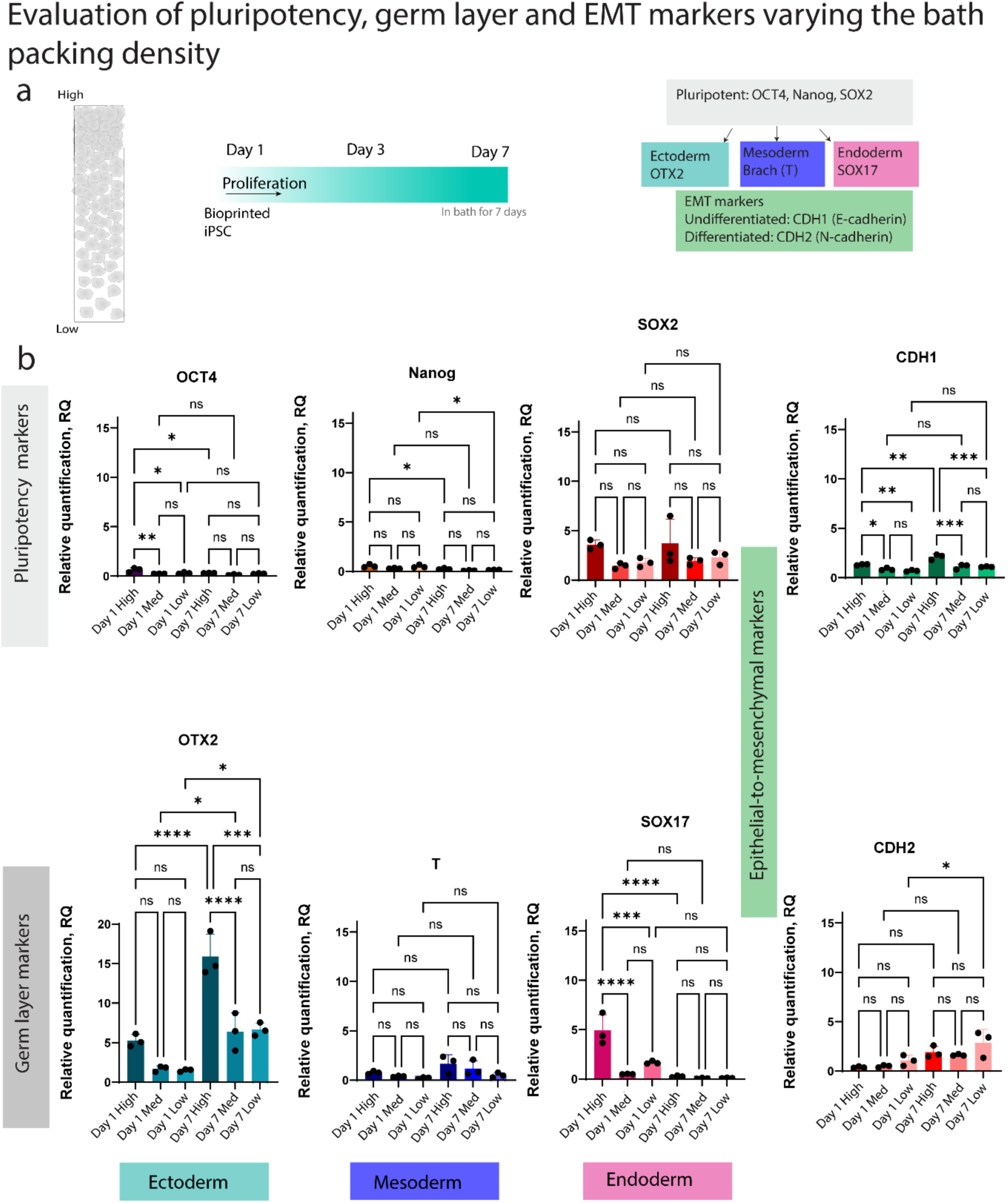
Pluripotency, germ layer, and EMT expression within bioprinted morphing iPSCs tissue constructs in a granular support bath: a) Schematic diagram representing the variation in packing density of the support bath (high, medium, and low) at the indicated time points selected for the evaluation of pluripotency, germ layer, and EMT markers using qRT-PCR. b) Plot of individual genes representing pluripotency (OCT4, Nanog, SOX2), germ layer (Ectoderm OTX2, Mesoderm T, and endoderm SOX17), and epithelial-mesenchymal transition (EMT) (E-cadherin CDH1 and N-cadherin CDH2) markers while the bioprinted tissue undergoes shape-morphing. Biological replicates n=3, one-way ANOVA was performed with Tukey’s multiple comparison test, where ns denotes not significant, * denotes p< 0.05, ** denotes p < 0.01, *** denotes p < 0.001, and **** denotes p < 0.0001.

**Supplementary figure 4.**
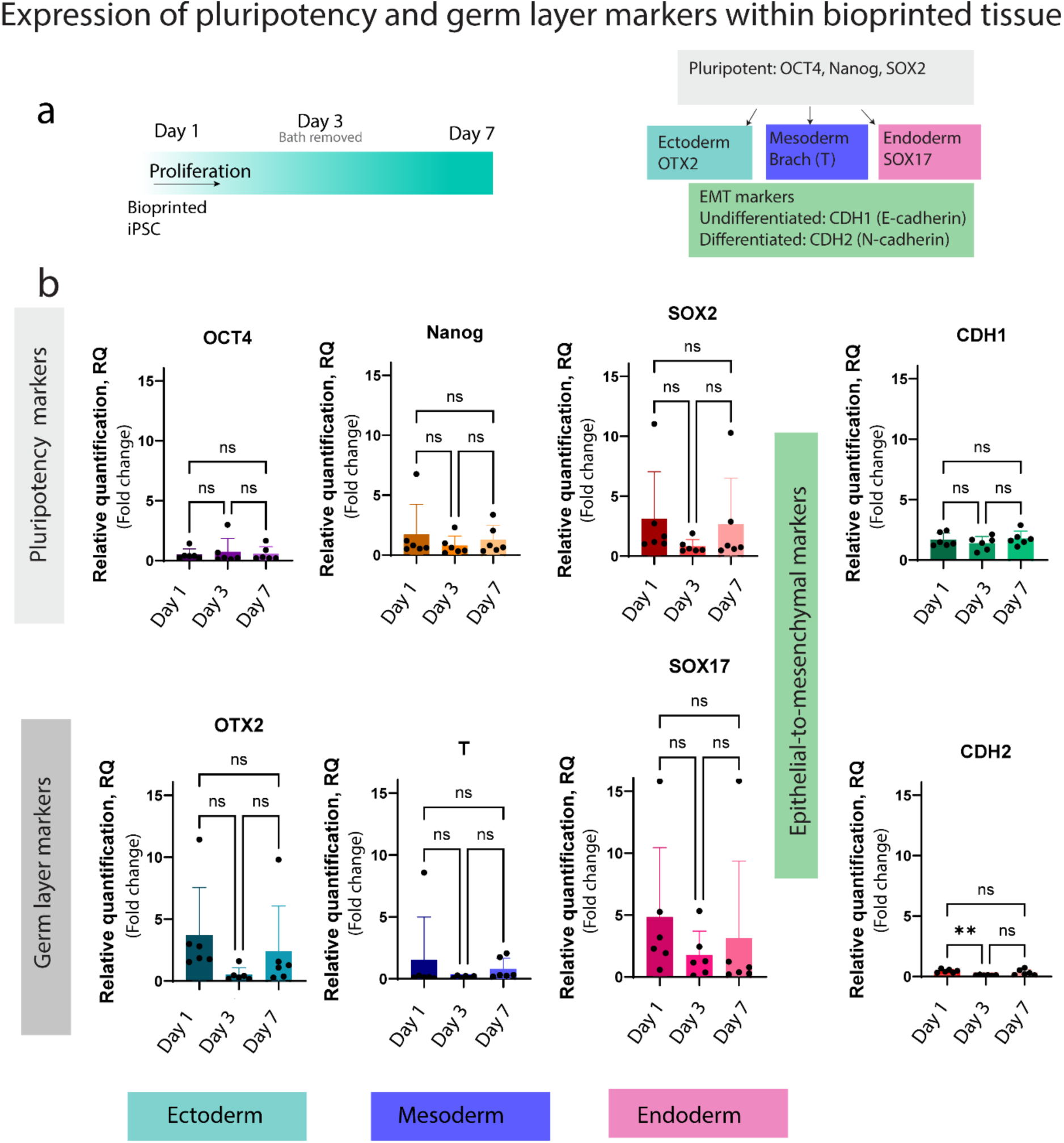
Expression of pluripotency, germ layer, and EMT within bioprinted morphing iPSCs tissue constructs: **a)** Schematic diagram indicating the time points selected for investigating the fold change of pluripotency, germ layer, and EMT expression (right). The panel of genes employed for qRT-PCR (left). **b)** Plot of individual gene representing pluripotency (OCT4, Nanog, SOX2), germ layer (Ectoderm (OTX2), Mesoderm (T) and endoderm (SOX17)) and EMT (E-cadherin CDH1 and N-cadherin CDH2) markers while the bioprinted tissue undergoes shape-morphing. Biological replicates n=6, one-way ANOVA performed with Tukey’s multiple comparison test, where ns denotes not significant, and ** denotes p < 0.01.

**Supplementary figure 5.**
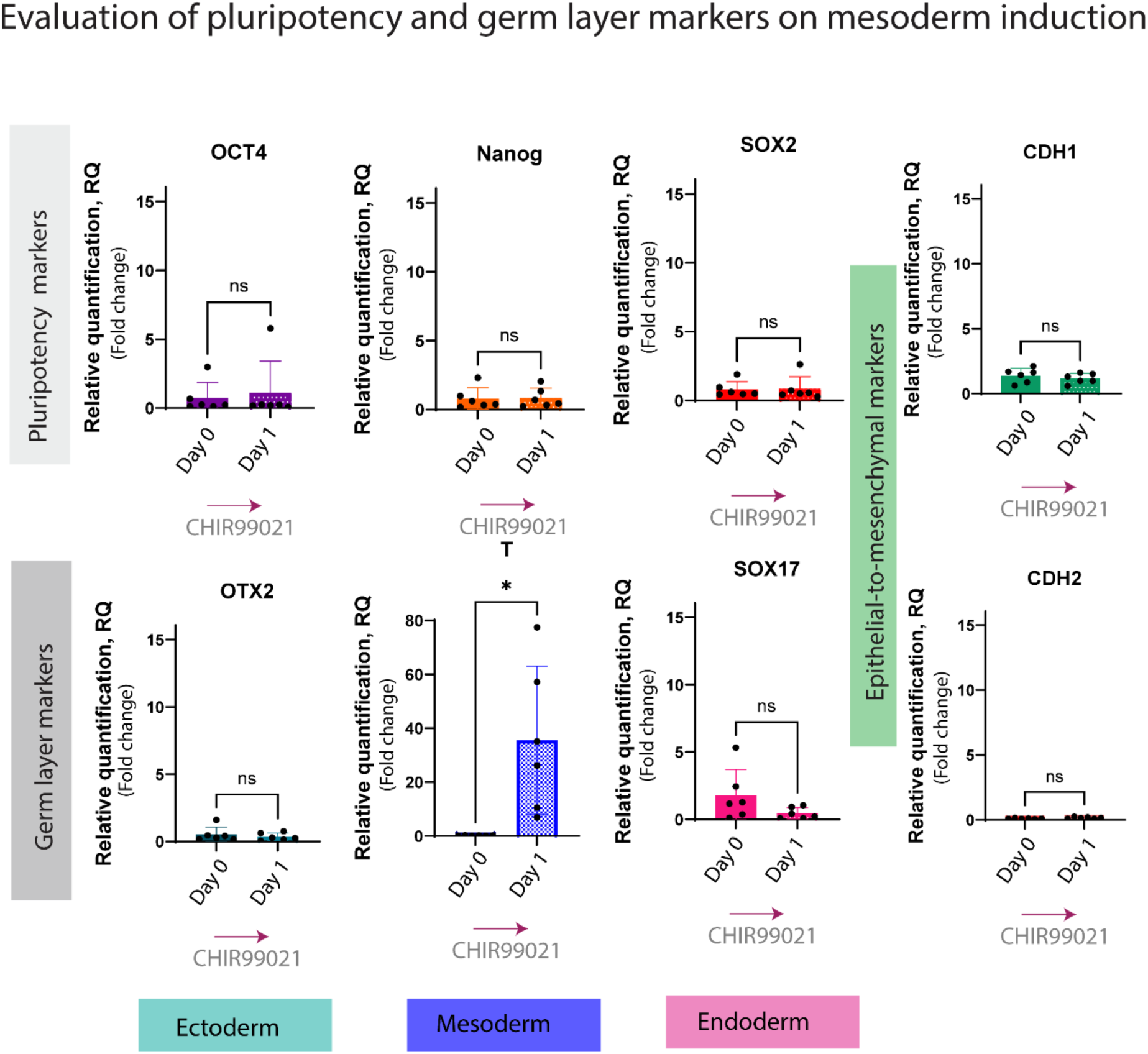
Evaluation of pluripotency, germ layer, and EMT expression within bioprinted constructs upon mesoderm induction by WNT activation: Plot of individual genes associated with pluripotency (OCT4, Nanog, SOX2), germ layer (Ectoderm OTX2, Mesoderm T, and endoderm SOX17), and EMT (E-cadherin CDH1 and N-cadherin CDH2) expression on mesoderm induction by activating the WNT pathway using CHIR99021, a small molecule that inhibits the GSK3 pathway and initiates mesoderm specification. Brachyury (T), which is related to mesoderm induction, showed significant upregulation after 24h post-CHIR99021 addition. All biological replicates (n = 6) and one-way ANOVA were performed with Tukey’s multiple comparison test, where ns denotes not significant, * denotes p < 0.05.

**Supplementary figure 6.**
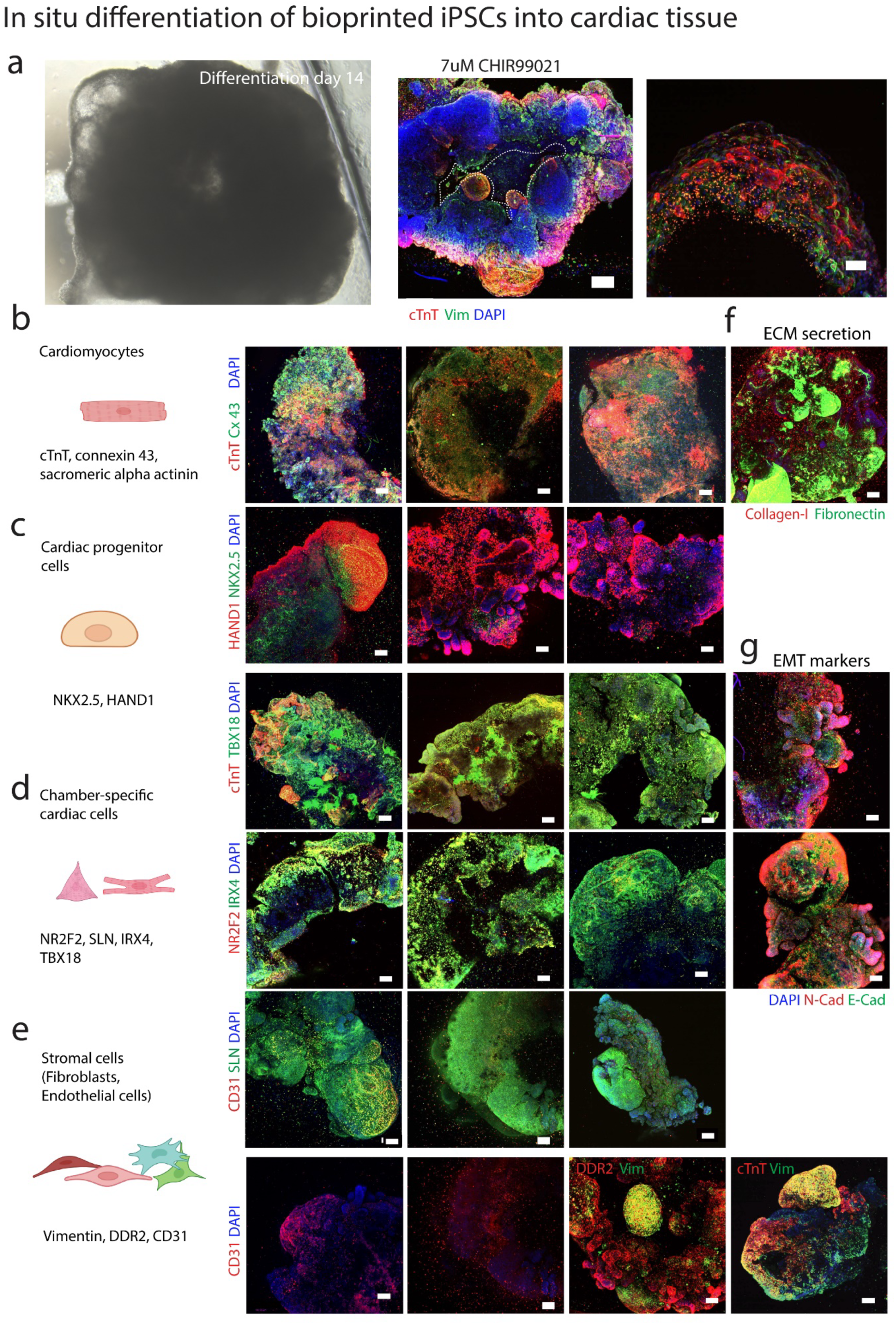
In situ differentiated multicellular bioprinted cardiac tissue: **a)** Brightfield image of in situ differentiated cardiac tissue using 6 µm CHIR99021 (left) and immunofluorescence images of differentiated tissues using 7 µm CHIR99021 (scale bars = 200 µm and 50 µm). Schematic representation (left) and immunofluorescence staining (right) of in situ differentiated cardiac tissues using 6 µm CHIR99021 for WNT activation related to **b)** cardiomyocyte markers (cTnT, cx43), **c)** cardiac progenitor cells marked by transcription factors (HAND1, KNX2.5), **d)** chamber-specific cardiac cell types: ventricular (IRX4) and atrial (NR2F2, SLN) and epicardium (TBX18), **e)** Stromal cell types such as fibroblasts and endothelial (DDR2, Vim, CD31) assessed using a panel of antibodies (scale 100 µm). **f)** Immunofluorescence images confirming ECM secretion (collagen-I, fibronectin), and **g)** EMT markers (E-cad, N-cad) within the constructs (Scale 100 µm). The lumens of the bioprinted ring constructs are indicated by white dashed lines.

## Supporting Information S3 – Additional Table

**Table S3.1:** Cell sources.

| Cell type | Catalogue no. | Company |
| --- | --- | --- |
| Gibco™ episomal hiPSC line | A18945 | Fisher Scientific |

**Table S3.2:**
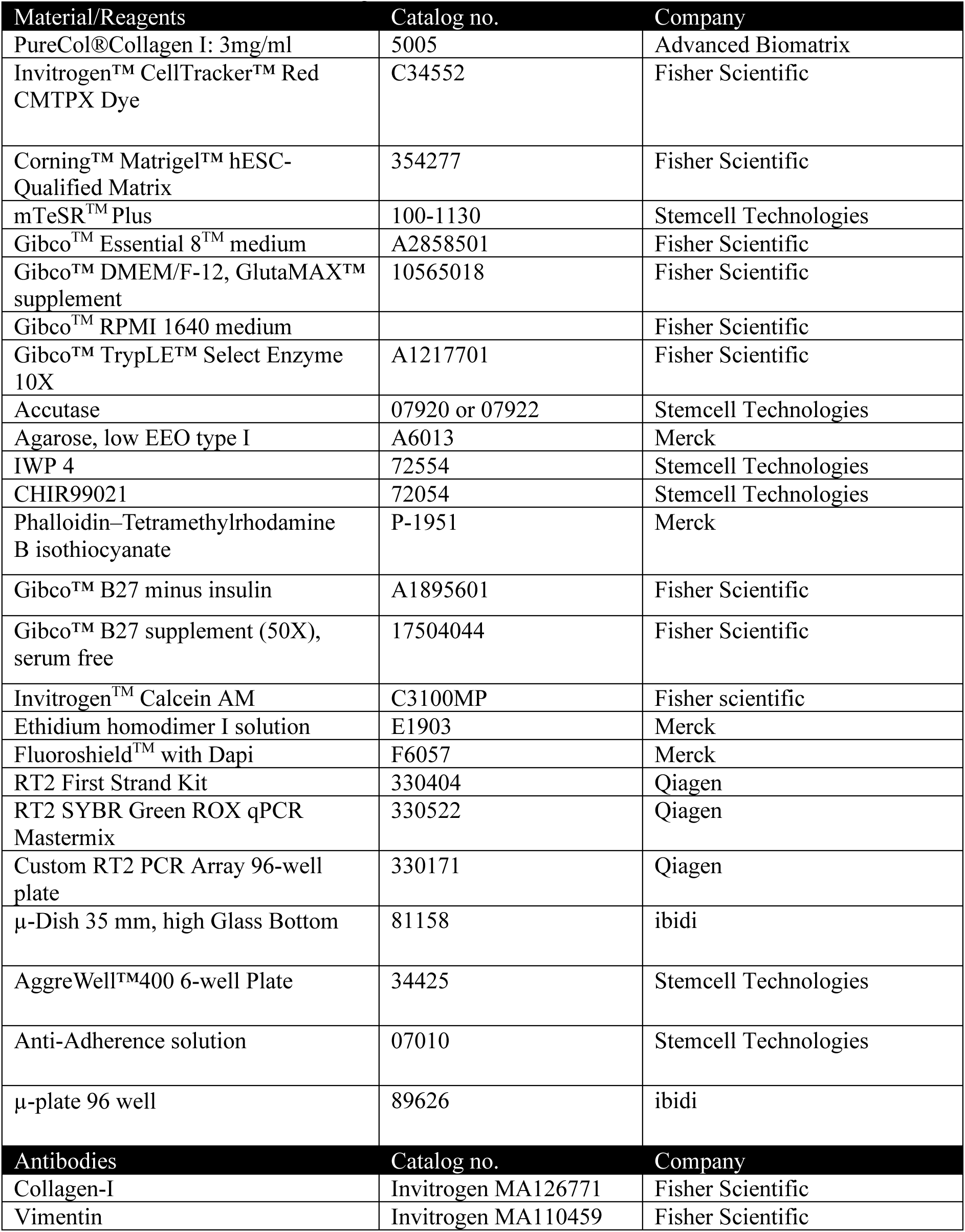
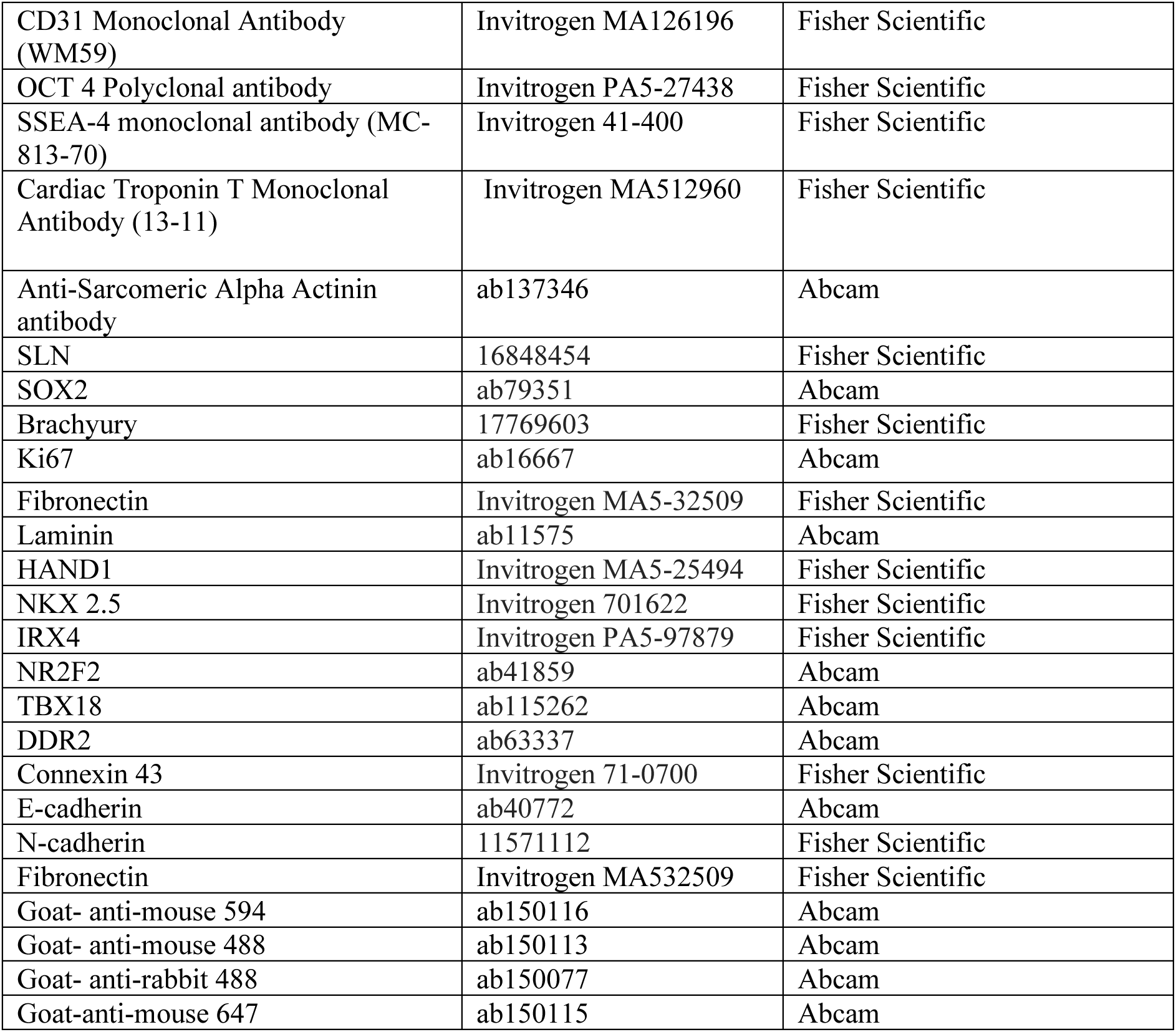
List of Materials and Reagents.

**Table S3.3:** Gene List for RT-PCR:

| Gene | Description code |
| --- | --- |
| **OCT4- Pluripotent** | Hs04260367_gH |
| **SOX2- Pluripotent** | Hs04234836_s1 |
| **Nanog- Pluripotent** | Hs02387400_g1 |
| **OTX2- Ectoderm** | Hs00222238_m1 |
| **T- Mesoderm** | Hs00610080_m1 |
| **SOX17-Endoderm** | Hs00751752_s1 |
| **CDH1-E-cadherin** | Hs01023895_m1 |
| **CDH2-N-cadherin** | Hs00983056_m1 |
| **Ki67- Cell Proliferation** | Hs01032443_m1 |
| **GAPDH- Housekeeping** | Hs99999905_m1 |

**Table S3.4:** Custom RT^2^ PCR array: Gene details.

| GenBank | Symbol | Description |
| --- | --- | --- |
| NM_003380 | VIM | Vimentin |
| NM_006475 | POSTN | Periostin |
| NM_001613 | ACTA2 | Actin, Alpha 2, smooth muscle, aorta |
| NM_006182 | DDR2 | Discoidin domain receptor tyrosine kinase 2 |
| NM_004387 | NKX2-5 | NK2 homeobox 5 |
| NM_002052 | GATA4 | GATA binding protein 4 |
| NM_003281 | TNNI1 | Troponin I type 1 (skeletal, slow) |
| NM_000364 | TNNT2 | Troponin T type 2 (cardiac) |
| NM_000363 | TNNI3 | Troponin I type 3 (cardiac) |
| NM_000165 | GJA1 | Gap junction protein, alpha 1, 43kDa |
| NM_181703 | GJA5 | Gap junction protein, alpha 5, 40kDa |
| NM_000891 | KCNJ2 | Potassium inwardly-rectifying channel, subfamily J, member 2 |
| NM_000719 | CACNA1C | Calcium channel, voltage-dependent, L type, alpha 1C subunit |
| NM_000335 | SCN5A | Sodium channel, voltage-gated, type V, alpha subunit |
| NM_001681 | ATP2A2 | ATPase, Ca ++ transporting, cardiac muscle, slow twitch 2 |
| NM_021223 | MYL7 | Myosin, light chain 7, regulatory |
| NM_000432 | MYL2 | Myosin, light chain 2, regulatory, cardiac, slow |
| NM_000257 | MYH7 | Myosin, heavy chain 7, cardiac muscle, beta |
| NM_002471 | MYH6 | Myosin, heavy chain 6, cardiac muscle, alpha |
| NM_002046 | GAPDH | Glyceraldehyde-3-phosphate dehydrogenase |

